# Effect of population structure and stabilizing selection on quantitative genetic variation

**DOI:** 10.64898/2026.03.29.714437

**Authors:** Juan Li, Joachim Hermisson, Himani Sachdeva

## Abstract

We study one of the simplest scenarios of polygenic selection that can be imagined: a subdivided population of diploid individuals expressing an additive trait under spatially homogeneous stabilizing selection. We are interested in the amounts of variation that can be maintained at mutation-selection-migration-drift equilibrium, at individual loci and at the level of the trait, within and among subpopulations. We derive analytical approximations for variance components and summary statistics such as *F*_*ST*_ and *Q*_*ST*_ under the assumptions of the infinite-island model and compare these with individual-based simulations. We find that: (i) There is a critical migration threshold (which depends on effect sizes of trait loci) below which population structure strongly inflates genic variance in the subdivided population to levels well above those in a panmictic population. Variation within each subpopulation is maximized close to the critical migration rate. (ii) The genetic basis of trait variation across subpopulations is most similar close to this migration threshold and (counter-intuitively) becomes less similar for higher migration rates. This has consequences for the ‘portability’ of Genome-Wide Association Studies (GWAS) between subpopulations, i.e, the extent to which loci with large contributions to variance in one subpopulation explain variance in other subpopulations. (iii) An analytical mean-field approach based on the single-locus diffusion approximation, together with effective migration and selection parameters (to account for associations between loci), very accurately predicts various quantities.

## Introduction

Large, outbreeding populations typically harbour high levels of molecular and phenotypic variation. However, we still do not fully understand how such variation emerges from the interplay between different evolutionary processes (Ch. 28, Walsh and Lynch (2018)). Early work focused on whether high trait variation might reflect a balance between mutation and stabilizing selection towards an intermediate trait optimum (Lande, 1975; Turelli, 1984; Bürger et al., 1989). More recent work suggests that a model of stabilizing selection acting on multiple traits is broadly consistent with the findings of Genome-Wide Association Studies (GWAS) for a suite of quantitative traits in humans (Simons et al., 2018, 2025).

This still leaves questions as to how much other factors, such as balancing or fluctuating selection and spatial population structure, contribute to the maintenance of variation, and whether observed patterns of variation might be consistent with several alternative evolutionary histories and scenarios (Barton and Keightley, 2002; Sella and Barton, 2019). Theoretical models show that selection that varies across time (Bürger and Gimelfarb, 2002; Bertram and Shafiei, 2025) or space (Slatkin, 1978; Barton, 1999; Tufto, 2000) can significantly inflate genetic variation under certain conditions. However, most of this work focuses on extreme cases, e.g., strongly selected loci (that evolve deterministically) vs. the infinitesimal model. This makes it difficult to generalise their conclusions to quantitative trait variation which may have a disproportionate contribution from loci of small to intermediate effect (Simons et al., 2018).

One general mechanism by which spatial population structure can increase genetic variability is if different subpopulations occupy different ‘adaptive peaks’, i.e., harbour distinct genetic combinations that are approximately equally fit (Wright, 1935). Alternative fitness peaks might arise due to spatially heterogeneous selection, e.g., if different trait values are favoured across different geographic regions or ecological niches. However, they can also arise under spatially uniform selection, provided that the genetic architecture of selected traits is sufficiently ‘redundant’, i.e., if many genetic combinations correspond to approximately the same phenotype (Goldstein and Holsinger, 1992; Phillips, 1996; Lythgoe, 1997; Barghi et al., 2020).

There has been much interest recently in how such genetic redundancy in adaptive traits could lead to (dis-)similar genomic signatures of adaptation in natural or experimental replicates (MacPherson and Nuismer, 2017; Yeaman et al., 2018; Barghi et al., 2019; Höllinger et al., 2023). However, all of this work assumes that replicate populations evolve independently. By contrast, most natural populations consist of sub-populations connected by historical or ongoing gene flow, which may cause the genetic basis of trait variation to become more similar across subpopulations, despite redundancy. This leads us to ask: do the same or different loci contribute to variation across different subpopulations? And how does this depend on selection and the genetic architecture of traits on the one hand, and dispersal and demography on the other? Put another way, to what extent does selection on polygenic traits perturb patterns of allele sharing at trait loci in a structured population, causing these to stand out from the neutral genomic background?

These questions are relevant to various efforts to predict phenotypes and make inferences about selection from genomic data while accounting for population structure. For instance, how well genomic prediction based on GWAS generalises across populations depends crucially on the extent to which the same genetic variants underlie phenotypic variation across populations with different ancestry (Duncan et al., 2019; Wang et al., 2020; Yair and Coop, 2022). Similarly, our ability to distinguish signatures of local vs. global adaptation in genomic data also depends on whether the same or different alleles underlie (either kind of) adaptation in different sub-populations (Ralph and Coop, 2015; Booker et al., 2021). However, even low levels of gene flow between populations can qualitatively alter theoretical expectations, which makes it crucial to model the combined effects of selection and population structure on trait variation.

As a step in this direction, we analyse the simplest possible scenario of a polygenic trait subject to spatially uniform stabilizing selection across a large number of ‘islands’ or sub-populations connected via migration. We explore the effects of selection and structure on diversity at individual trait loci as well on the total trait variance, which also depends on statistical associations (or linkage disequilibria; LD) between trait loci. The goal is to describe and predict patterns of variation at different ‘scales’: globally (across the full population) vs. locally (within sub-populations), and at the locus vs. the trait level.

Previous work on the effect of (uniform) selection and spatial structure on quantitative genetic variation is largely simulation-based (Goldstein and Holsinger, 1992; Latta, 1998; Le Corre and Kremer, 2003, 2012; McDonald and Yeaman, 2018) or, otherwise, derives analytical results by assuming strong selection and deterministic evolution (Phillips, 1996; Lythgoe, 1997). This makes it difficult to link theory to patterns in genomic variation, which are significantly affected by genetic drift. Here, we address this gap by developing detailed analytical theory that accounts for the combined effects of mutation, stabilizing selection, drift and population structure on trait variants, thus allowing for a more comprehensive understanding. Our analysis relies on the fact that stabilizing selection on a polygenic trait primarily generates selection against variance (at least if the trait mean is close to the trait optimum). This generates selection against the minor allele at any trait locus, resulting in the same evolutionary dynamics as at a locus with underdominance(Wright, 1935; Robertson, 1956; Bulmer, 1972). Thus, as a first approximation, we can treat the polygenic trait as being determined by a set of *independently* evolving underdominant loci. This allows us to use single-locus theory, in particular, the diffusion approximation to study the stochastic evolution of allele frequencies at trait loci (Wright, 1937; Bulmer, 1972; Barton and Rouhani, 1993). We then build upon this to account for the effects of LD between trait loci using effective parameters. As we demonstrate below, this approach yields accurate predictions for gene-level as well as trait-level variation across most of parameter space, and also extends to traits influenced by a broad distribution of effect sizes.

A natural reference point for our analysis is the strong migration limit of the model, in which the subdivided population resembles a very large, panmictic population. The total genetic variance at mutation-selection equilibrium in such a population depends only on the total mutation rate for the trait, and the strength of stabilizing selection (Latter, 1960; Bulmer, 1972), and is independent of the effect size of trait loci, at least in the House-of-Cards mutation regime (Turelli, 1984). How might this change in scenarios with limited migration between subpopulations? In particular, do loci with certain effect sizes contribute disproportionately to variability within or between subpopulations, making them more likely to drive adaptive response and/or be identified in genomic data?

## Model and Methods

### Model

We consider a subdivided population consisting of *D* demes (islands), each with *N* diploid individuals. For theoretical analyses, we will take the limit *D* → ∞, which corresponds to the infinite-island model, introduced by Wright (1931). Demes are connected via migration: in each generation, a migrant pool is formed by drawing individuals from across the whole population, with migration probability *m* per individual. Individuals from this pool are then uniformly redistributed back across all demes of the population. Thus, *Nm* represents the average number of migrants exchanged between a deme and the rest of the population.

Each individual expresses an additive trait influenced by *L* unlinked (freely recombining), bi-allelic loci. Alternative alleles at any locus are denoted by *X* = 0, 1; the genotype of an individual *j* at locus *i* is represented as 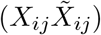, where *X*_*ij*_ and 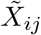 denote maternally and paternally inherited alleles. Thus, there are three possible genotypes: (00), (01) (which is equivalent to (10)), and (11) at locus *i*, with contributions to trait value given by ™*α*_*i*_, 0, and *α*_*i*_ respectively. Hereafter, we refer to *α*_*i*_ as the effect size at locus *i*. The trait value *z*_*j*_ of an individual *j* is given by:

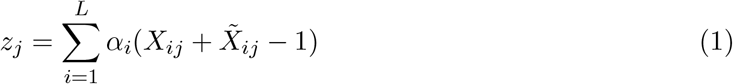

For simplicity, we assume that the trait has no environmental component. However, this assumption is not crucial, since a non-zero environmental variance only rescales the effective strength of stabilizing selection on the trait (see, e.g., Bürger et al., 1989). Polymorphism is maintained by mutation, which occurs at rate *µ* per allele per individual per generation. Thus, the expected number of trait-affecting mutations per diploid individual per generation, is 2*µL*.

The trait is assumed to be under stabilizing selection, which disfavours individuals with trait values that deviate from the optimum. The fitness of an individual with trait value *z* is:

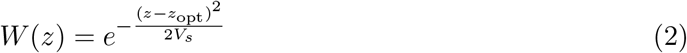

The trait optimum, *z*_opt_, and the width of the fitness function, 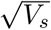, is the *same* across all demes, resulting in spatially *uniform* selection across the entire population.

Finally, we assume that individuals mate randomly within demes based on the Wright-Fisher model. Thus, the next generation is formed by sampling 2*N* parents (with replacement) within each deme, with sampling weights proportional to the relative fitness of individuals in that deme. Each of these 2*N* parents produce haploid gametes via free recombination between trait loci. Gametes are then paired randomly to form *N* diploid offspring, which replace the parents.

All three processes – stabilizing selection, migration, and drift – can generate LD between trait loci within demes, thus causing them to evolve in a non-independent manner. If recombination is much faster than other evolutionary processes, such LD is weak, and can be neglected to a first approximation. This is the basis of our linkage equilibrium (LE) approximation, which accounts for the effects of migration, mutation, genetic drift and direct selection (arising from its own effect) at each trait locus, but not those of LD between loci. However, LD cannot always be ignored, particularly with intermediate levels of population structure, and if traits have a large total mutation rate 2*µL*. Therefore, we extend our approximations to account for the effects of LD using appropriately defined effective parameters, which leads to more accurate predictions for the variance contributions of trait loci. Finally, we develop predictions for the variance of trait values, and how it is partitioned within and between demes.

### Allele frequency distributions at trait loci under Linkage Equilibrium (LE)

Following Bulmer (1972), we describe the evolution of allele frequencies at trait loci in continuous time. To approximate the dynamics of the discrete generation model via continuous time equations, we will assume that the distribution *ρ*_*j*_(*z*) of trait values within any (say, the *j*^*th*^) deme is normal with mean 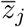 and variance *V*_*W,j*_, i.e.,

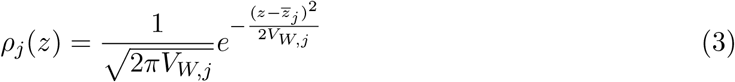

Here, *V*_*W,j*_ is the *within*-deme phenotypic variance, which is equal to the genetic variance (since trait values are assumed to have no environmental component).

The mean fitness in the deme is then given by:

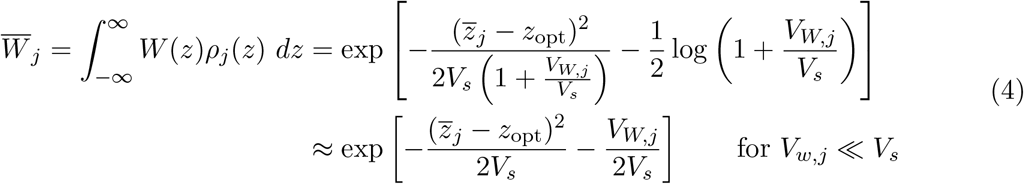

Thus, for *V*_*W*_ ≪ *V*_*s*_, the reduction in (log) mean fitness has two distinct components – one depending on 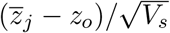, the deviation of the trait mean from the trait optimum, and the other on 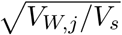, the standard deviation of trait values, both expressed relative to the width of the stabilizing selection function.

The trait mean can be expressed in terms of the allele frequencies as 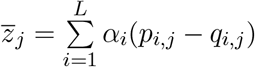 where *p*_*i,j*_ and *q*_*i,j*_ = 1 ™ *p*_*i,j*_ denote, respectively, the frequencies of the trait-increasing and trait-decreasing alleles at locus *i* in deme *j*. The genetic variance, *V*_*W*_, has two components: a genic component, 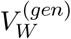, and a component due to LD, 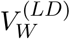. If recombination between trait loci is much stronger than selection and gene flow, individual demes are approximately in linkage equilibrium (LE), so that: 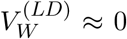, and 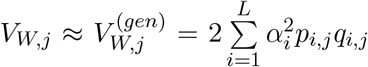. Then, the selection gradient at trait locus *i* in deme *j* is simply:

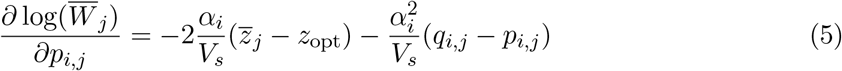

Note that the same expression for the selection gradient can be obtained without assuming a normal distribution of trait values, if we start with a model of continuous time evolution and assume LE (Höllinger et al., 2023).

The trait mean within any deme rapidly approaches the trait optimum, at least if there is no mutational bias and the optimum is close to the centre of the phenotypic range, i.e., *z*_opt_ ≈ 0 (see also Bulmer, 1971). Thus, in a population at equilibrium, the selection gradient at any locus is approximately 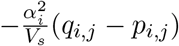. Furthermore, if mutation, migration and selection per locus are sufficiently weak, i.e., if *µ, m*, 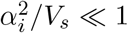, then we can describe the deterministic evolution of allele frequencies (neglecting drift) in continuous time as follows:

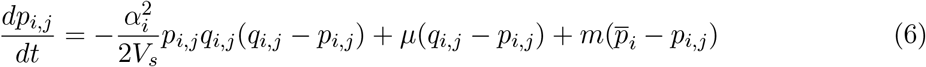

The first term describes the effects of selection on allele frequencies, and has exactly the same form as for a single underdominant locus with heterozygote disadvantage 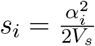. The second term describes the effects of mutation, which tends to drive allele frequencies towards 1*/*2 (in the case of symmetric mutation). The third term describes the effects of migration that pulls allele frequencies towards 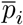, the mean allele frequency (at locus *i*) in the migrant pool, which is equal to the mean allele frequency across the entire population under the infinite-island assumption.

Thus, allele frequencies at any locus underlying a trait under stabilizing selection in a subdivided population follow relatively simple dynamics, which are identical to those of a single underdominant locus. As discussed above, this holds if: first, genetic variance within demes is sufficiently small, i.e., *V*_*W*_ */V*_*S*_ ≪ 1; second, LD between trait loci is negligible; third, the trait mean in each deme is close to the trait optimum, i.e., 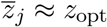. In subsequent sections, we will relax the first and second assumption.

Equation (6) describes the deterministic dynamics of allele frequencies in a large population under LE. However, genetic drift generates additional random fluctuations, with the variance of random allele frequency change per generation due to drift being equal to: Var[Δ*p*_*i,j*_] = *p*_*i,j*_*q*_*i,j*_*/*(2*N*) within any deme. We can describe the combined effects of drift, migration, mutation and selection on allele frequencies using the diffusion approximation (Kimura, 1964). In particular, under the infinite-island model (i.e., in the limit *D* → ∞), allele frequencies {*p*_*i,j*_} across different demes *j* = 1, … *D* (at say, locus *i*) are mutually *independent* random variables that depend only on the mean allele frequency 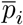 in the migrant pool. From eq. (6), it follows that the equilibrium distribution of *p*_*i,j*_ in any deme *j*, conditioned on 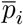 is:

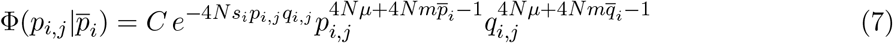

where *C* is the normalization constant (see also Wright, 1937). In the limit *D* → ∞, the variance of 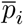 approaches zero. Thus, the allele frequency in the migrant pool, 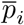, must be *deterministically* equal to the expected allele frequency within demes. In other words, we have:

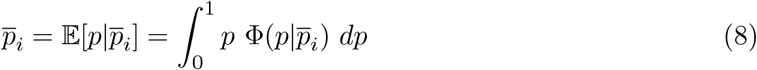

This can be solved numerically (see also Barton and Rouhani, 1993), thus allowing us to calculate various other quantities by integrating over eq. (7). We refer to the predictions obtained in this way as LE predictions. In particular, one can express 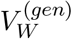 and 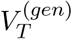, the expected *genic* variance within demes and across the total population as:

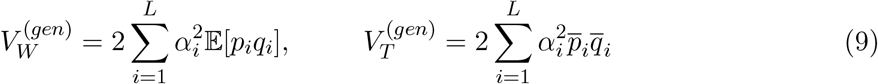

Here 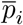 and 𝔼[*p*_*i*_*q*_*i*_] can be obtained from eq. (7) via numerical integration. In addition, we derive simple analytical expressions for these quantities in the weak selection (4*Nµ < Ns* ≲ 1) and strong selection (*Nµ* ≪ 1 ≲ *Ns*) limits (Sec. SI.a, SI).

In the following, we will report trait variances in units of *V*_*\**_, the genic variance in an infinitely large panmictic population at mutation-selection equilibrium. Setting *m* = 0 in eq. (6), it follows that at mutation-selection equilibrium, we have *p*_*i*_*q*_*i*_ = *µ/s*_*i*_, provided 4*µ < s*_*i*_. This gives: *V*_*\**_ = 4*µLV*_*s*_, which is proportional to the total mutation rate for the trait and independent of effect sizes (Turelli, 1984).

Thus, the *scaled* genic variances can be expressed as:

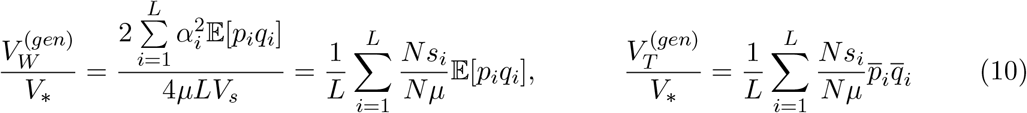

Since 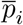 and 𝔼 [*p*_*i*_*q*_*i*_] only depend on the *Ns, Nµ* and *Nm* (eq. (7)), this allows us to express our results purely in terms of scaled (dimensionless) parameters.

#### Distribution of variance contributions of trait loci

In addition to predicting the expected genic variance, the equilibrium allele frequency distribution 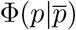 (eq. (7)) can also be used to predict the *distribution* of variance contributions of individual loci within demes. As before, we scale the variance contribution *v* of any locus by the deterministic expectation *v*_*\**_(which is 4*µV*_*s*_ for a single locus), and denote the scaled variance by 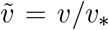. A locus with effect size *α* (or selection coefficient 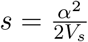) and allele frequency *p* within a deme has a (scaled) variance contribution 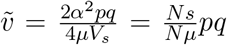. Transforming variables from *p* to 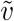 in eq. (7) gives the following expression for the the probability density 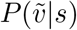 of variance contributions:

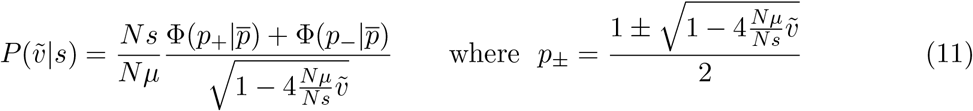

The cumulative probability that a trait locus with selective effect *s* contributes an amount *exceeding y* (in units of *v*_*\**_) to the genic variance within a deme is then:

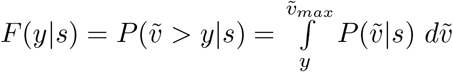

where 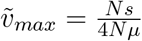 is the maximum possible value of 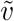 (corresponding to *p* = 1*/*2).

Similarly, we can define 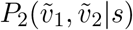 as the joint probability density of variance contributions of a locus with selective effect *s* in two demes (arbitrarily labeled ‘1’ and ‘2’). The corresponding joint cumulative distribution is: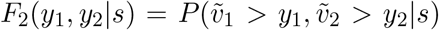. Since allele frequencies in different demes are independent under the infinite-island model (conditional on the mean 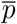), we have: 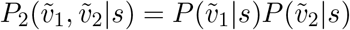 and *F*_2_(*y*_1_, *y*_2_|*s*) = *F* (*y*_1_|*s*)*F* (*y*_2_|*s*).

In general, trait loci will have unequal effects, leading to a distribution of selective effects, which we denote by *ψ*[*s*]. The cumulative distributions *F* (*y*) and *F*_2_(*y*_1_, *y*_2_) across all trait loci can then be obtained by integrating over *ψ*[*s*]:

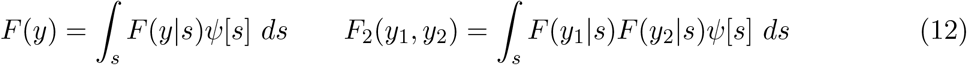

### Accounting for LD using effective parameters

So far, we have neglected the effects of LD between trait loci within demes. However, both stabilizing selection and migration build up LD, leading to non-independent evolution of trait loci. In this section, we will discuss each of these two sources of LD, and describe how the effects of LD on a given locus can be encapsulated via a suitably defined effective selection coefficient and effective migration rate. These effective rates depend on allele frequencies at all other trait loci, allowing us to obtain a mean field (i.e., self-consistent) solution for all the allele frequencies jointly. This approach of representing the effects of LD using effective parameters is very similar in spirit to the Quasi Linkage Equilibrium (QLE) approximation and applies as long as the dynamics of LD are much faster than the dynamics of allele frequencies (Kimura, 1965). This, in turn, requires recombination to be much faster than all other processes that influence single-locus dynamics, which is the case in a model with unlinked loci.

First, consider the effects of stabilizing selection, which generates negative LD between trait-increasing (or, equivalently, trait-decreasing) alleles within a deme. This causes the genetic variance (i.e., the variance of trait values) to be less than the genic variance (Bulmer, 1971). To quantify this reduction, we assume (as before) that *L* is large enough that the distribution *ρ*(*z*) of trait values within any deme is approximately normal with mean 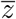 and variance *V*_*W*_. Stabilizing selection changes the distribution of trait values to: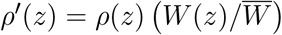, where 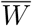 is given by eq. (4). This shifts the trait mean towards the trait optimum and decreases the trait variance, so that after selection, we have:

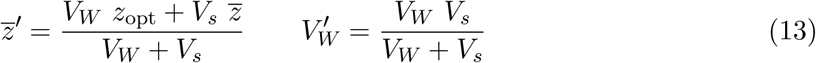

Random mating within demes leaves the trait mean unchanged 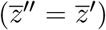, but changes the trait variance to: 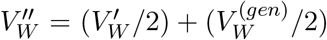. Here, 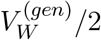 is the segregation variance (also known as the within-family variance) released by recombination between parental haplotypes. For a randomly mating population, the segregation variance is equal to half of the genic variance. For a population at equilibrium, the trait value distributions do not change from one generation to the next. Thus, we have: 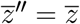 and 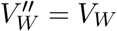, which gives:

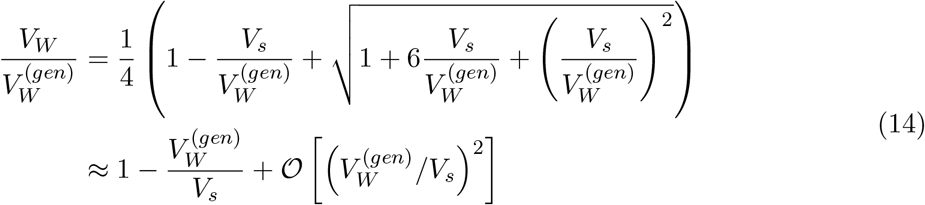

which is equivalent to eqs. 18 and 22 of Bulmer, 1971. Thus, for 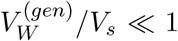, the genetic variance within a deme is reduced relative to genic variance by an amount 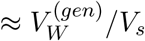, which is twice the genetic load per deme.

In addition to reducing genetic variance, negative LD between trait loci also generates indirect selection at each locus by causing the trait-increasing allele at any locus to become associated with alleles of opposite effect at other loci. This slightly reduces the average effect of the allele (e.g., as estimated by GWAS), which slightly relaxes heterozygote disadvantage and increases heterozygosity at each locus, thus causing the genic variance to be *higher* than the LE prediction. Further, in parameter regimes where negative LD is significant (i.e., when 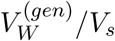 is non-negligible), the approximation 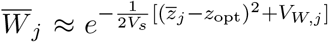 (second line of eq. (4)) also becomes less accurate. Instead, we must use the full expression in eq. (4) to calculate the (direct) selection gradient any locus. Following Veller and Coop (2024) and Negm and Veller (2024) (see eq. (25) in the latter), we account for both these effects by assuming that each trait locus still evolves as if under underdominant selection, but with an effective selection coefficient:

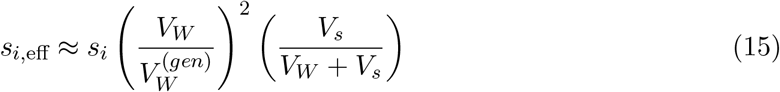

The factor 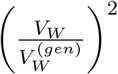 captures the effect of negative LD between trait loci, while the factor 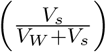 corrects for the effect of non-negligible *V*_*W*_ */V*_*s*_ on direct selection at the trait locus. From eq. (14), it follows that *s*_*i*,eff_*/s*_*i*_ depends only on 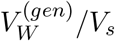.

Next, consider how migration contributes to LD between trait loci. Since all demes are adapted to the same optimum, migrants are phenotypically very similar to residents but may bear distinct genetic combinations, especially at low levels of migration, when different demes carry alternative minor alleles. Mating between migrants and residents produces F1 hybrids which are phenotypically similar to both parents but can have much higher heterozygosity. Descendants of these F1s (e.g., back-cross individuals) thus tend to be phenotypically more variable and have lower average fitness than the descendants of residents. This reduces the longterm contribution of migrant individuals to the gene pool and generates indirect selection against immigrant alleles due to their association with less fit genetic backgrounds (which deviate more than average from the trait optimum). Following (Sachdeva, 2022) and (Surendranadh and Sachdeva, 2025), we account for this by assuming that the effect of migration on allele frequencies is still described by eqs. (6) and (7), but with *m* replaced by an effective migration rate *m*_eff_. In the SI, we show that under the assumptions of our model:

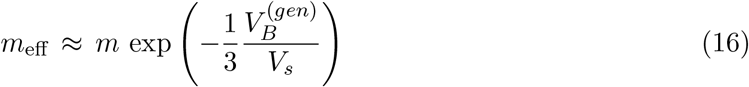

where 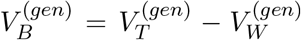, is the among-deme genic variance (see eq. S.24). Thus, the reduction in effective migration rates is strongest when 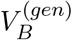 is highest, i.e., when demes are nearly fixed for alternative alleles at trait loci. See also Veller and Simons, (2024) for a related derivation of *m*_eff_ in a two-deme setting.

We now have a system of equations for the effective parameters {*s*_*i*,eff_} and *m*_eff_, and the variance components 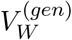 and 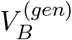, which can be solved self-consistently. Technically, this is best done using an iterative process, where we start with an initial ‘guess’ for 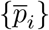, then obtain the various genic variances by integrating over 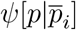, and substitute these in eqs. (14)-(16) to compute {*s*_*i*,eff_} and *m*_eff_. Finally, we substitute these values of {*s*_*i*,eff_} for *s*_*i*_ and *m*_eff_ for *m* in eq. (7) to solve for the new 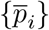. These steps are iterated until {*s*_*i*,eff_}, 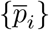 and *m*_eff_ converge to fixed values. For the various cases that we have tested, this iterative procedure converges to a unique solution (for {*s*_*i*,eff_}, 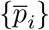 and *m*_eff_) regardless of the choice of initial 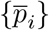, even though convergence is not guaranteed in theory. We refer to the predictions obtained in this way as LD predictions, since they capture the effect of LD via effective parameters.

### Variance among demes and total variance across the full population

So far, we have focused on the genic and genetic variance within demes. In this section, we develop analytical approximations for *V*_*B*_, the genetic variance among demes, which is defined as: 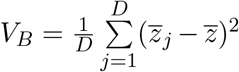, where 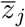 is the trait mean in the *j*^*th*^ deme and 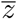 the trait mean across the entire population. This allows us to also calculate the total genetic variance *V*_*T*_ = *V*_*W*_ + *V*_*B*_.

Calculating *V*_*B*_ requires us to first find 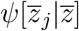, the distribution of the trait mean in any (say, the *j*^*th*^) deme, conditional on the trait mean across the full population. Following Barton and Rouhani (1993), the expected rate of change of the trait mean in deme *j* can be written as:

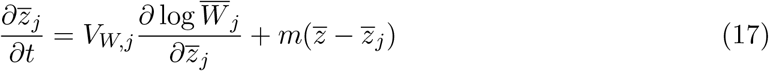

The first term describes the effects of selection on the trait mean, under the assumption that the trait value distribution within any deme has a negligible third moment (skew). The second term describes the effect of migration, which tends to pull the trait mean within any deme towards the population average. Equation (17) neglects the effect of mutation on trait means, which is justified as long as there is no mutational bias, and the trait optimum is at the centre of the phenotypic range.

Using eq. (17), together with the fact that mean squared fluctuations in trait mean per generation (due to genetic drift) go as *V*_*W,j*_*/N*, we can write down the following expression for 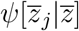 under the diffusion approximation (see also Barton and Rouhani, 1993):

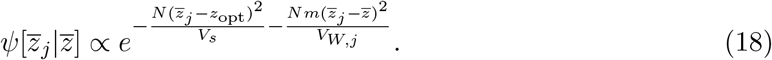

If the number of demes is large enough that fluctuations in 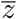 are negligible (so that 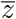 is deterministically equal to *z*_opt_), then the trait mean within any deme is normally distributed with mean 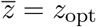 and variance *V*_*B*_ given by:

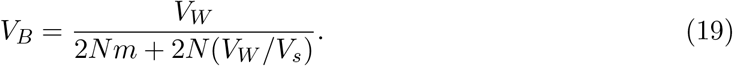

Note that in the limit of no migration (i.e., as *Nm* → 0), eq. (19) reduces to *V*_*B*_ = *V*_*s*_*/*(2*N*), which is the predicted variance of the trait mean in a single population subject to stabilizing selection (Lande, 1976). In the absence of selection, (i.e., as *V*_*s*_ → ∞), eq. (19) reduces to: 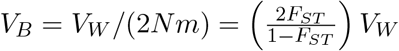, which is the neutral expectation (Whitlock, 1999).

Equation (19) can now be used to predict *Q*_*ST*_, which is defined as the ratio of the among-deme variance to the total genetic variance expected in a hypothetical panmictic population with the same allele frequencies as the subdivided population (Spitze, 1993). *Q*_*ST*_ is a measure of quantitative genetic differentiation between populations, and in our model is given by:

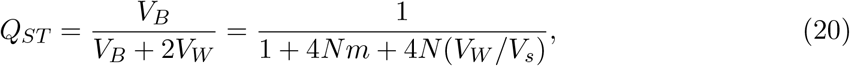

We can also use our approximations to predict the total genetic variance across the full population using *V*_*T*_ = *V*_*W*_ + *V*_*B*_. In general, the total genetic variance *V*_*T*_ will differ from the genic variance 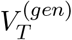 because of LD between trait loci as well as deviations from Hardy-Weinberg equilibrium (across the population as a whole). Thus, comparing *V*_*T*_ and 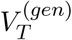 provides an estimate of the combined strength of intra and inter-locus statistical associations, which, in turn, reflect the combined effects of population structure and stabilizing selection. For the special case of equal-effect trait loci, we have: 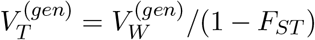, where 1 ™ *F*_*ST*_ is measured as the ratio of the average within-deme heterozygosity averaged across all trait loci to the average total heterozygosity (see also Weir and Cockerham, 1984). This gives:

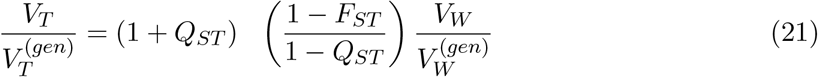

Note that for a neutrally evolving trait, *Q*_*ST*_ = *F*_*ST*_, and 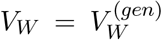, so that 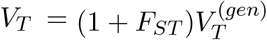 (Wright, 1949; Whitlock, 1999). Thus, neutral population structure *increases* genetic variance relative to genic variance by creating a deficit of heterozygotes across the population as a whole, which causes per-locus contributions to *V*_*T*_ to be more extreme than one would expect in a panmictic population with the same allele frequencies. By contrast, stabilizing selection *reduces* genetic variance relative to genic variance by ‘locking in’ alleles with opposite effects on trait value, thus creating negative LD between trait loci and reducing *V*_*W*_ and (especially) *V*_*B*_. Equation (21) describes how these two opposing effects together shape genetic variance. In particular, it suggests that genetic variance should be significantly lower than genic variance (i.e., the effect of negative LD between trait loci should dominate), provided *F*_*ST*_ at trait loci is high and *Q*_*ST*_ is low (see also Le Corre and Kremer, 2003).

The arguments outlined in this section (and in deriving eq. (14) in the previous section) are framed purely in terms of trait means and variances. We thus refer to eqs. (14), (19) and (20) as quantitative genetic (QG) predictions. As before, we can build upon these to account for the effects of LD within demes using effective parameters. This is accomplished by replacing *m* by *m*_eff_ in eqs. (19) and (20), and by using *m*_eff_ and {*s*_*i*,eff_} in eq. (7) when computing 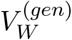 (which then predicts *V*_*W*_). We refer to these as *QG*(eff) predictions.

### Simulations

We test analytical predictions by comparing against discrete-generation, individual-based simulations which track multi-locus genotypes of all diploid individuals across all demes. In each generation, the life cycle consists of mutation, followed by selection and mating within demes, which is then followed by migration. We perform two kinds of simulations: the first with underdominant selection on a single locus (Fig. 1), and the second with stabilizing selection on an additive trait influenced by *L* loci, where effect sizes at trait loci are either all equal (as in Fig. 2) or drawn from a distribution (Fig. 3). See also the Model description above.

**Figure 1.**
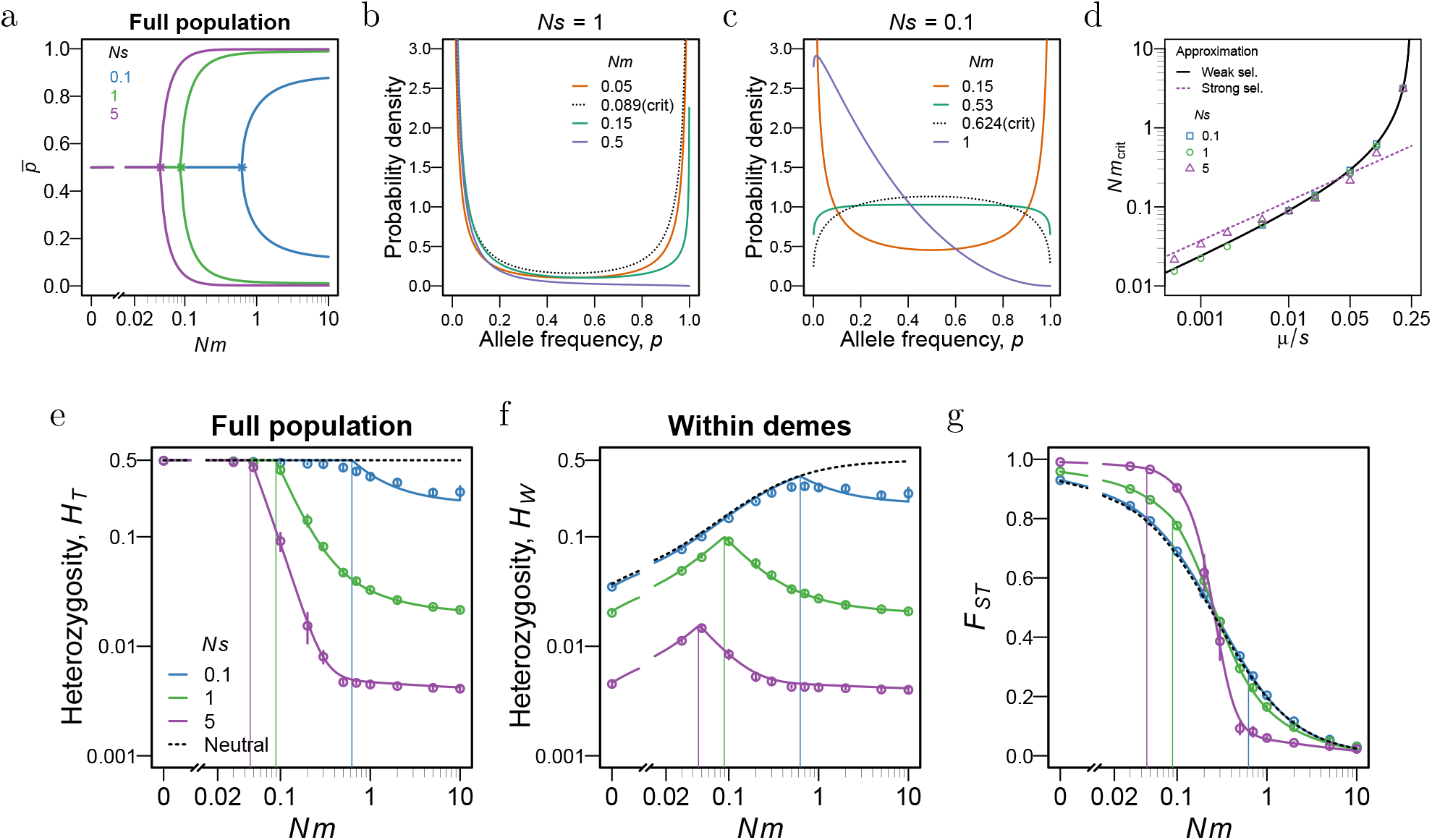
Genetic diversity at a single underdominant locus under the island model. (a) Mean allele frequency across the full population, 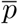, as a function of *Nm* for different *Ns* values (colors), at equilibrium. There is a critical migration threshold *Nm*_crit_ (marked by a *), such that there is only one stable equilibrium 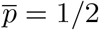 for *Nm < Nm*_crit_, but two alternative equilibria 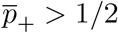 and 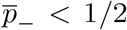 for *Nm > Nm*_crit_. (b)-(c) Equilibrium probability density, *ψ*[*p*], of the allele frequency *p* in a single deme, for different values of *Nm* (colors), including *Nm*_crit_ (dashed black curve), for *Nµ* = 0.01 and scaled selection coefficient (b) *Ns* = 1 or (c) *Ns* = 0.1. The probability density is symmetric around *p* = 1*/*2 if *Nm < Nm*_crit_ but biased towards *p* (which we assume to be equal to 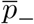) if *Nm > Nm*_crit_. All solid lines in (a)-(c) are obtained from the diffusion approximation (eq. 7). (d) The scaled critical migration rate *Nm*_crit_ as a function of *µ/s* for *Nµ* = 0.01, and different values of *Ns*. Symbols show *Nm*_crit_ as obtained from the diffusion approximation; the solid line shows the ‘weak selection’ approximation for *Nm*_crit_ (eq. S.4) which is independent of *Ns* (for a given *µ/s*); the dashed line shows the ‘strong selection’ approximation for *Nm*_crit_ (eq. S.11) for *Ns* = 5. (e) Total heterozygosity across the full population, 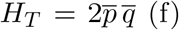 Within-deme heterozygosity, *H*_*W*_ = 𝔼[2*pq*] and (g) expected *F*_*ST*_ at selected locus as a function of *Nm*, for different values of *Ns*, and *Nµ* = 0.01. Solid lines show predictions of the diffusion approximation (eq. 7); dashed lines represents the neutral prediction, and circles show results of individual-based simulations with *D* = 100 demes and *N* = 200 diploids per deme, where each simulation is run for 5 × 10^5^ generations. Each circle shows the mean obtained by averaging over 10 simulation replicates and over the last 10^4^ generations (at intervals of 10^3^ generations) within each replicate; error bars show 1.96 times the standard error (SE). For reference, we also indicate *Nm*_crit_ thresholds (as predicted by the diffusion approximation) using vertical lines in panels (e), (f) and (g).

**Figure 2.**
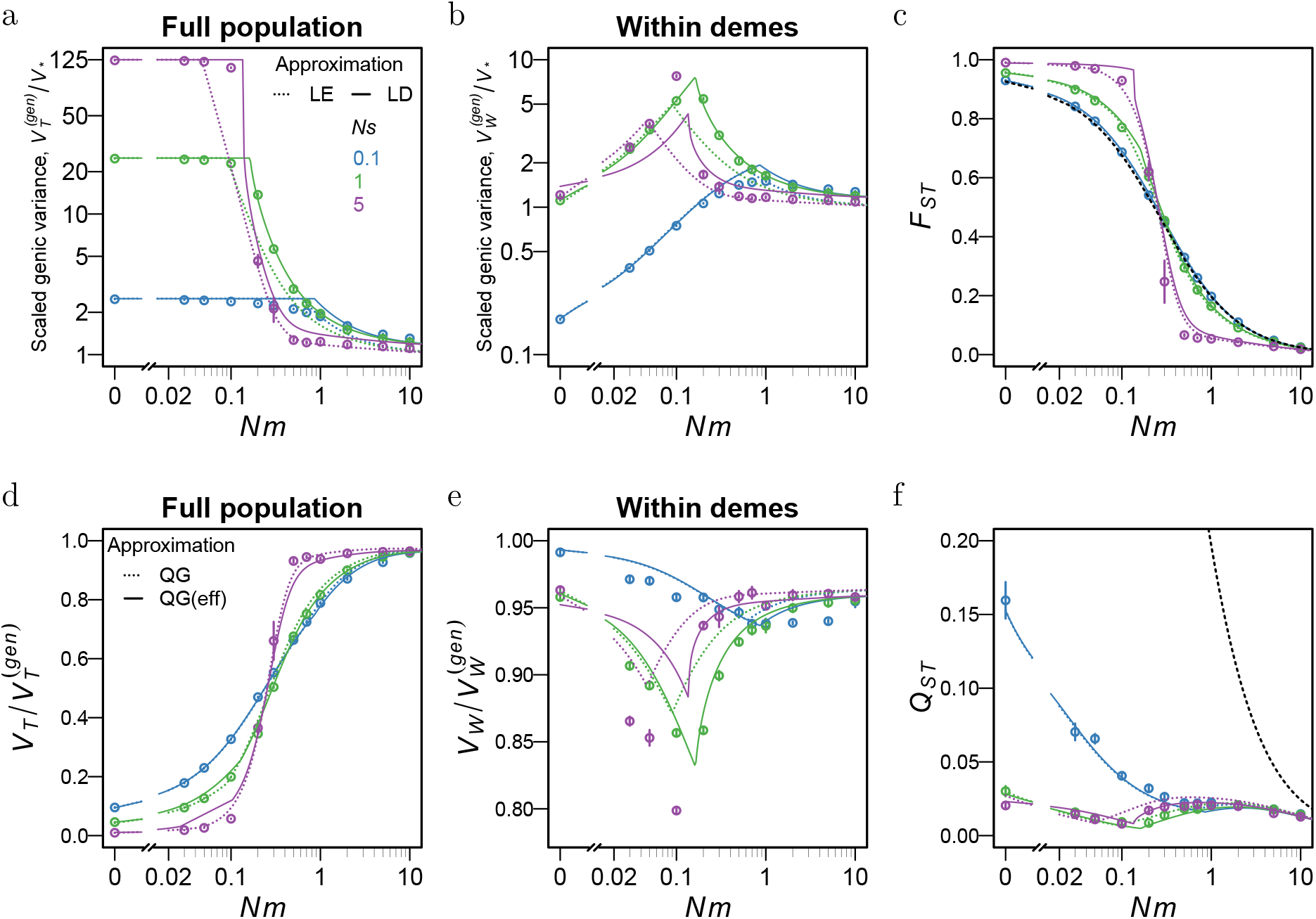
Effect of stabilizing selection and population structure on genic variance, genetic variance, *F*_*ST*_ and *Q*_*ST*_, for an additive trait influenced by equal-effect loci. The expected (a) total genic variance across the full population, 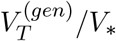, and (b) within-deme genic variance, 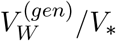, as a function of *Nm*, the average number of migrants per deme per generation. Genic variances are scaled by 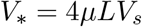, the variance in a large panmictic population at deterministic mutation-selection balance (neglecting LD). (c) Average *F*_*ST*_ across all trait loci vs. *Nm*, (d) Ratio of the genetic variance to the genic variance in the population as a whole,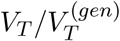, and (e) the corresponding ratio within demes, 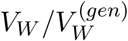, as a function of *Nm*. (f) *Q*_*ST*_ vs. *Nm*. The different colors show results for different values of *Ns*, the scaled selection coefficient per locus, where 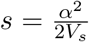 and *a* is the effect size of the locus. For a trait with equal-effect loci, the scaled variances can be expressed as: 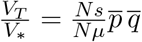 and 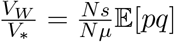. Symbols show results of individual-based simulations with *D* = 100 demes and *N* = 200 diploids per deme that express an additive trait influenced by *L* = 200 equal-effect loci, with scaled mutation rate *Nµ* = 0.01 per locus. Each circle shows the mean obtained by averaging over 10 simulation replicates and over the last 10^4^ generations (at intervals of 5 × 10^3^ generations) within each replicate; error bars show 1.96 times the standard error. Simulations are run for 10^5^ generations, except those with *Ns* = 5 and *Nm* ≤ 0.3, which are run longer (for 5 × 10^5^ generations). The dashed lines in Figs. 2(a)-(c) show LE predictions based on the diffusion approximation for an underdominant locus (and are the same as the predictions in Figs. 1(e)-(g) up to a constant scaling factor). Solid lines in Figs. 2(a)-(c) show LD predictions that account for the effect of LD on heterozygosity at trait loci using effective parameters. Dashed lines in 2(d)-(f) show QG predictions based on eqs. (14), (19) and (21) together with single-locus theory for the genic variances; solid lines show QG_eff_ predictions that use effective parameters.

**Figure 3.**
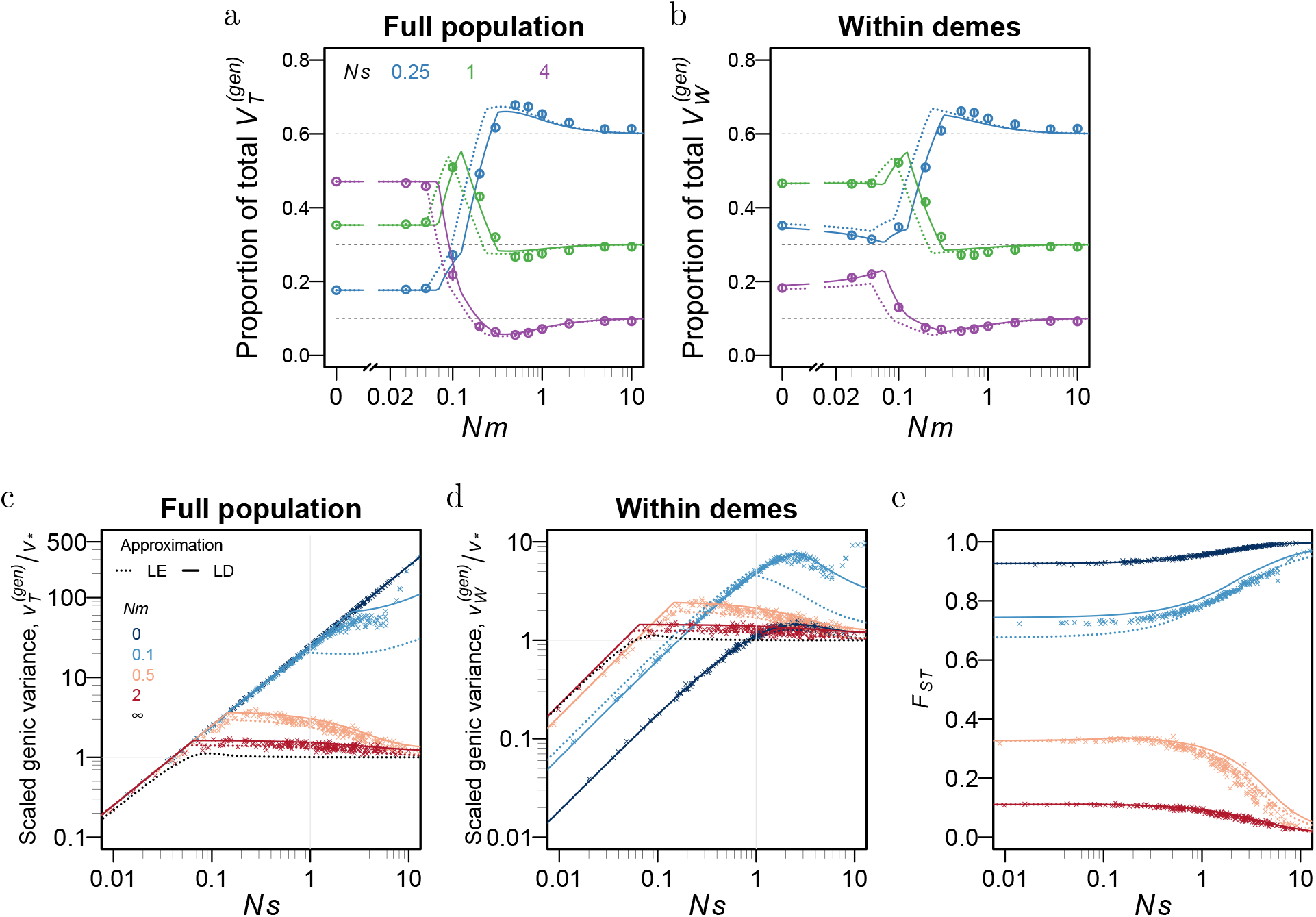
Variation in traits influenced by loci of unequal effect. (a)-(b) Proportion of (a) total genic variance and (b) within-deme genic variance contributed by small-effect (*Ns* = 0.25), medium-effect (*Ns* = 1) and large-effect (*Ns* = 4) loci, for a trait influenced by *L* = 200 unlinked loci, of which 60%, 30% and 10% have small, medium and large effects respectively. The three colors correspond to the three different effect sizes. (c)-(e) Expected contribution of each trait locus to (c) within-deme genic variance and (d) total genic variance across the full population, and (e) expected *F*_*ST*_ at each trait locus, as a function of the scaled selection coefficient (*Ns*) of the locus, for a trait influenced by *L* = 200 loci with effect sizes drawn from an exponential distribution with 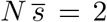. Here, per-locus variance contributions are scaled by the deterministic expectation, *v*_*\**_ = 4*µV*_*s*_. Different colors correspond to different migration rates; dashed black line shows LE prediction in the limit *Nm* → ∞, i.e., for a single *panmictic* population with population size *N*_tot_ = *ND*. Circles (in (a)-(b)) and points in (c)-(e)) show results of individual-based simulations with *D* = 100 demes and *N* = 200 diploid individuals per deme, with scaled mutation rate *Nµ* = 0.01 per locus. Each circle/point shows the mean obtained by averaging over 10 simulation replicates and over the last 10^4^ generations (at intervals of 5 × 10^3^ generations) within each replicate. The dashed lines show LE predictions based on the diffusion approximation for an underdominant locus; solid lines show LD predictions using effective parameters. In (c)-(e) and for *Nm* = 0.1, loci with the largest *Ns* values have not yet equilibrated in simulations run for 5 × 10^5^ generations and initialised with allele frequencies at each locus in each deme drawn independently from a beta distribution *Beta*[4*Nµ*, 4*Nµ*].

Simulations are performed with two different kinds of initial conditions – we either draw allele frequencies at each locus *in each deme* independently from a beta distribution *Beta*[4*Nµ*, 4*Nµ*], or first draw the mean allele frequencies 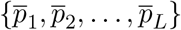 at each locus from the beta distribution and then set the allele frequencies in individual demes to be equal to the respective 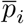. As shown in Fig. S5, SI, the time taken for the population to reach equilibrium can be quite different, depending on the initial conditions (especially at low migration rates), even though the equilibrium state itself is independent of initial conditions. Simulations are run for *D* = 100 demes and *N* = 200 individuals per deme, and a maximum of 10^6^ generations, which is found to be sufficient for equilibration in all cases (including very weak migration), with equilibration being faster at larger migration rates (see also Fig. S5).

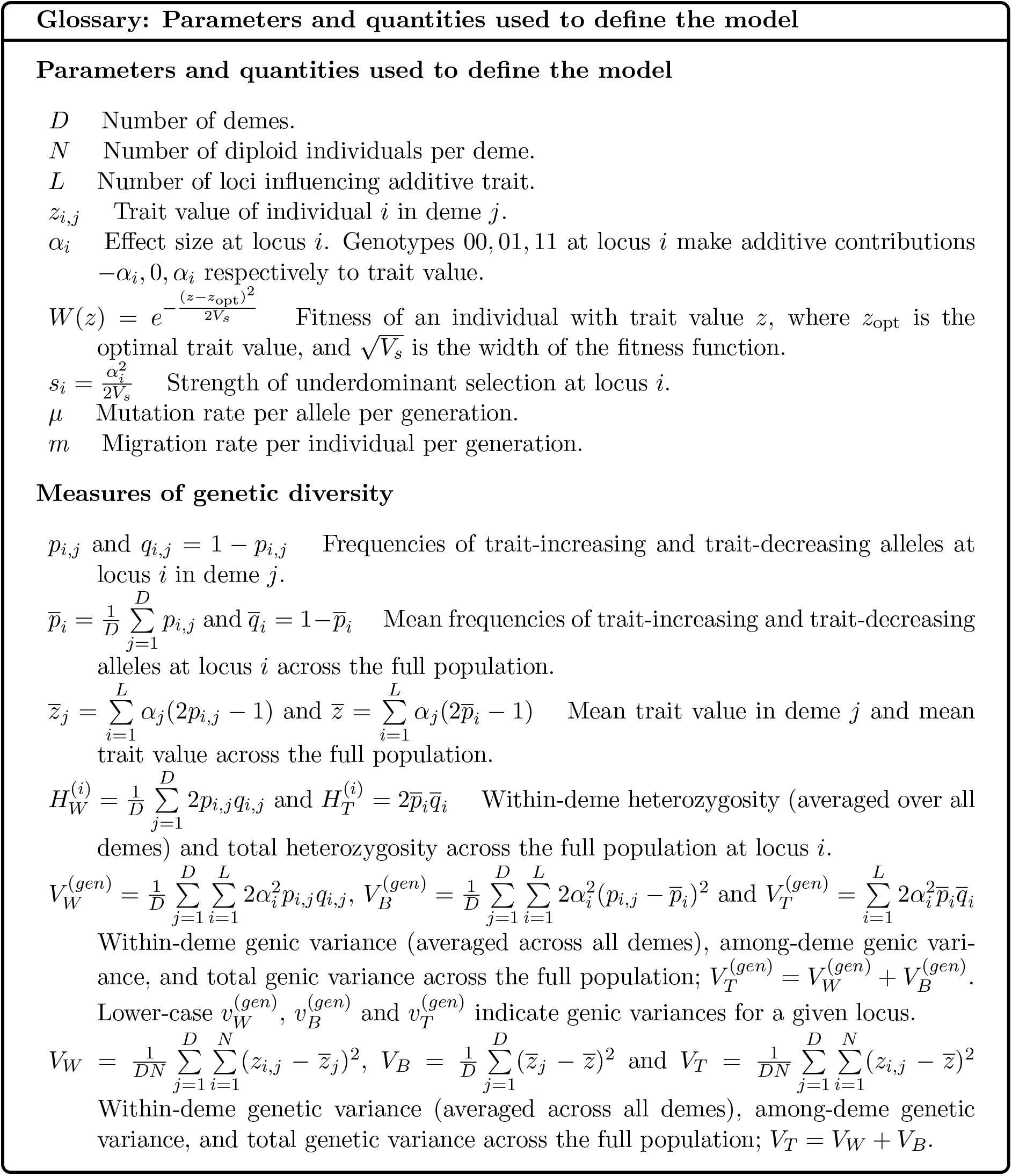
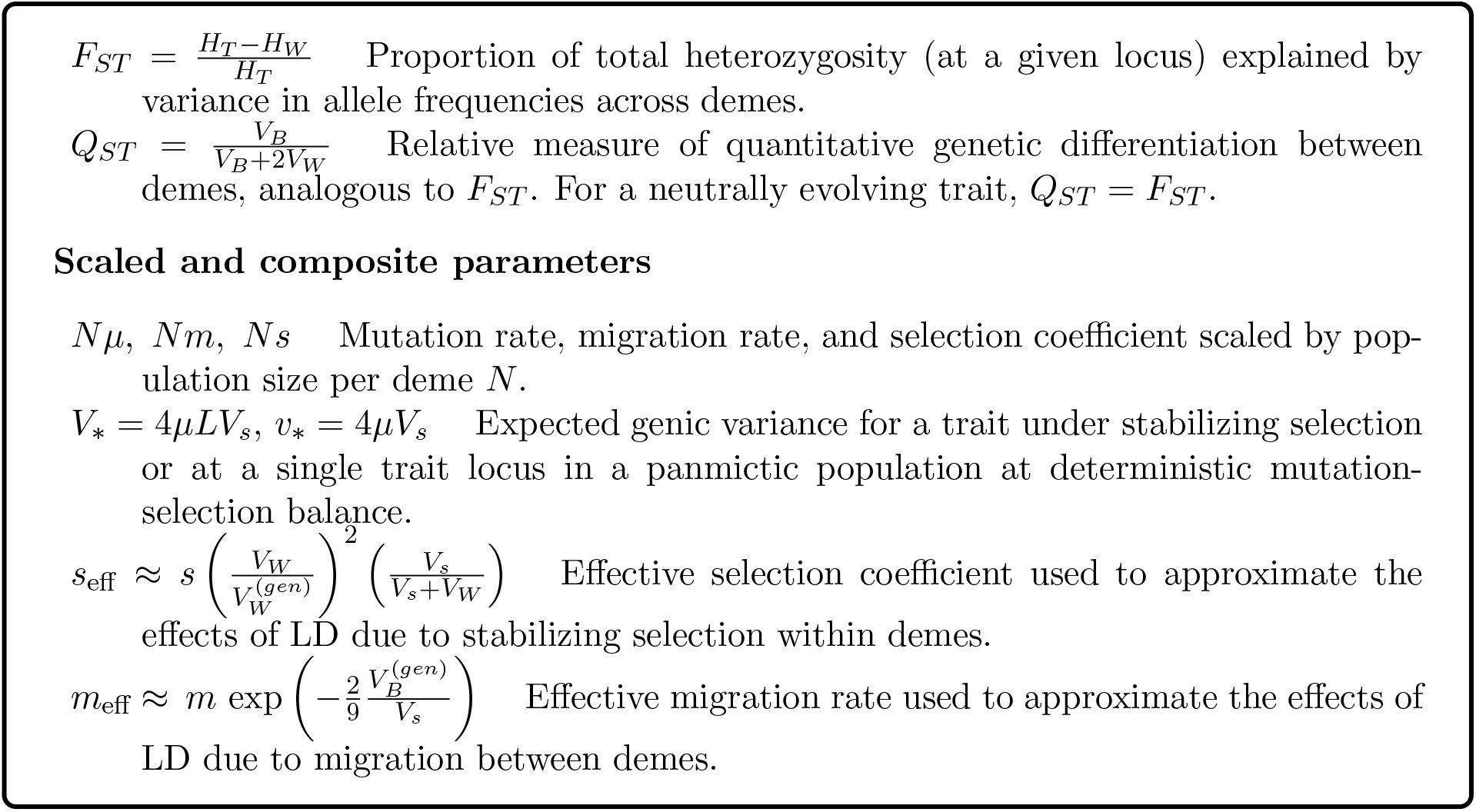

## Results

As discussed above, stabilizing selection on a polygenic trait essentially manifests as under-dominant selection on individual trait loci when the trait mean is at the trait optimum. Thus, as a precursor to analysing the polygenic model, we first consider the behaviour of a single locus under underdominant selection in the infinite-island setting. We then build upon this to investigate how variation in the polygenic trait – at individual loci (genic variance) and in trait values (genetic variance) – is partitioned within and among demes. This requires us to go beyond single-locus theory and also account for LD between trait loci, which we do via effective selection and migration parameters (see Methods). Finally, we investigate questions of direct empirical relevance: how many and what kind of loci make atypically large contributions to trait variance, and to what extent such outliers are shared across demes. Throughout, we compare the results of individual-based simulations with LE and LD predictions, thus clarifying to what extent LD influences patterns of variation and whether we can describe its effects succinctly in terms of a handful of effective parameters.

### Single locus under underdominant selection

Consider a single biallelic locus under underdominant selection (Figure 1). If selection, migration and mutation are sufficiently weak (*s, m, µ* ≪ 1), and if the population size per deme is large (*N* ≳ 50), then the allele frequency distribution at migration-mutation-selection-drift equilibrium depends only on the scaled parameters *Ns, Nm* and *Nµ* via eq. (7). The population exhibits two qualitatively different types of equilibrium states, separated by a critical (scaled) migration rate *Nm*_crit_ (Fig. 1a; see also Barton and Rouhani,1993).

At low migration rates (i.e., for *Nm* ≤ *Nm*_crit_), alternative alleles increase to high frequencies in different demes, leading to high levels of polymorphism in the population as a whole. The mean allele frequency across the full population is 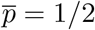, and the total population-wide heterozygosity is 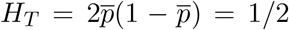, regardless of *Nm* (Fig. 1e). The allele frequency distribution within demes is symmetric around 1*/*2 (Figures 1b, 1c); it is U-shaped if selection is strong, i.e., *Nm < min*(1*/*2, *Nm*_crit_) (neglecting 𝒪(*Nµ*) terms), but inverted U-shaped if selection is weak, i.e., 1*/*2 *< Nm < Nm*_crit_ (see, e.g., green curve in fig. 1c).

By contrast, for *Nm > Nm*_crit_, the evolution of different demes is strongly coupled, and the *same* allele reaches high frequency in the majority of demes. The mean frequency *p* across the full population evolves towards one of two possible equilibrium values, 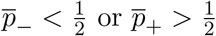. The total heterozygosity *H*_*T*_ is thus less than 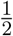, and decreases with increasing migration, approaching 2(*µ/s*) (the expected heterozygosity in a large panmictic population) at large values of *Nm* (Fig. 1e). The allele frequency distribution within demes is no longer symmetric about 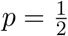. Instead, it is unimodal and concentrated around 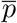 if 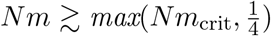 or U-shaped with a strong skew towards 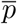 if 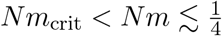 (see, e.g., green curve in fig. 1b).

In Section SI.a, SI, we derive approximate expressions for the critical migration rate and show that *Nm*_crit_ depends primarily on *µ/s*, and follows:

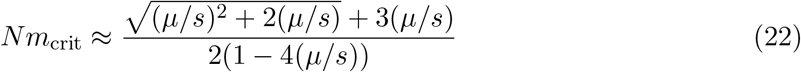

for *Ns* ≲ 1 and *µ/s <* 1*/*4 (solid black line in fig. 1d). Thus, as selection becomes stronger (and *µ/s* smaller), lower levels of migration are sufficient to couple evolution across demes. Note that we restrict ourselves to *µ/s <* 1*/*4, since at higher mutation rates, underdominant selection is overwhelmed by mutation, driving allele frequencies to *p* = 1*/*2.

We also track *H*_*T*_ and *H*_*W*_, the total and within-deme heterozygosity, as well as *F*_*ST*_ at the selected locus, and compare analytical predictions based on eq. (7) with individual-based simulations of *D* = 100 demes and *N* = 200 individuals per deme (Figs. 1e-1g). Simulations are conducted for three different selection strengths (*Ns* = 0.1, 1, 5) – representing weak, intermediate, and strong selection relative to local drift. The corresponding predictions for *Nm*_crit_ are 0.624, 0.089 and 0.046 (depicted by vertical lines in Figs. 1e and 1f). These figures show close agreement between simulations and theory. Slight deviations for weak selection and *Nm* ≃ *Nm*_crit_ are due to the relatively modest total population size *N*_*tot*_ = *ND* (see Section SI.c), and vanish as the number of demes, and consequently *N*_*tot*_, increases (Figure S2).

Figures 1e and 1f shows that the effect of migration on heterozygosities differs qualitatively depending on whether *Nm* is above or below the critical migration threshold *Nm*_crit_. As described above, both *H*_*T*_ and *H*_*W*_ decrease with increasing migration for *Nm > Nm*_crit_, but are unchanged (in the case of *H*_*T*_) or increase (in the case of *H*_*W*_) with *Nm*, if *Nm < Nm*_crit_. In particular, *H*_*W*_, the heterozygosity within demes, is maximised at *Nm* ~ *Nm*_crit_, being lower at both weaker and stronger levels of migration – when either individual demes or the population as a whole are close to fixing one of the alleles. Thus, while strong population subdivision allows high genetic variation to be maintained across the full population, intermediate levels of subdivision are most effective at maintaining genetic variation within local demes.

Interestingly, *Nm*_crit_ is not associated with any qualitative change in the behaviour of *F*_*ST*_, which decreases smoothly with increasing migration (Figure 1g). For *Nm < Nm*_crit_, this decrease is purely due to the increase in *H*_*W*_ (since *H*_*T*_ is equal to 1*/*2, regardless of *Nm*). By contrast, for *Nm > Nm*_crit_, the decrease in *F*_*ST*_ primarily reflects changes in *H*_*T*_, which decreases much more sharply than *H*_*W*_ with increasing *Nm* (compare, e.g., figures 1e and 1f).

Figure 1g also shows how the strength of underdominant selection influences *F*_*ST*_. Under weak selection (*Ns* = 0.1), *F*_*ST*_ is almost indistinguishable from the neutral expectation (dashed black line), regardless of *Nm*. In contrast, for stronger selection (*Ns* = 1 or 5), *F*_*ST*_ at the selected locus is higher than neutral *F*_*ST*_ for 4*Nm* ≲ 1, but then drops below it for 4*Nm* ≳ 1. To better understand these observations, we develop simpler analytical approximations (described in Sec. SI.a, SI) for the weak selection (*Nµ* ≪ *Ns* ≲ 1) and strong selection (*Nµ* ≪ 1 ≲ *Ns*) limits. These show that *F*_*ST*_ goes from slightly above the neutral prediction to slightly below it at 4*Nm* = 1 + 𝒪 (*µ/s*) for any locus with 4*Nm*_crit_ *<* 1, which translates into *µ/s* ≲ 1*/*24 (see eq. (22)), i.e., moderate or strong underdominant selection.

Interestingly, for such loci, the shape of the allele frequency distribution also changes qualitatively – from asymmetric U-shaped (a minority of demes close to fixing the alternative allele) to single-peaked (same allele common across all demes) – at a migration threshold 4*Nm* = 1 + 𝒪 (*µ/s*). This threshold is very close (but not exactly equal) to the threshold at which *F*_*ST*_ falls below the neutral prediction. Indeed, the two thresholds coincide as *µ/s* → 0. More generally, while increasing selection reduces both *H*_*W*_ and *H*_*T*_, its effect on the total heterozygosity, *H*_*T*_, is weaker if 4*Nm* ≲ 1, i.e., if at least some demes remain locally fixed for the alternative allele. Thus, in this regime, *F*_*ST*_ increases with increasing selection.

### Polygenic trait under stabilizing selection

Building upon the single-locus analysis of the previous section, we now ask: how much variation is maintained in a polygenic trait under stabilizing selection towards a uniform optimum? Stabilizing selection has two main effects – it pushes the trait mean towards the trait optimum, and it reduces the trait variance. The first effect is typically quite strong, causing 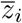, the trait mean within any deme *i*, to change as: 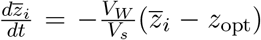, where *V*_*W*_ is the within-deme trait variance. Thus, trait means tend to rapidly approach the trait optimum, so that 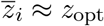 in expectation, and 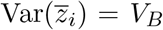 is small (≲ *V*_*s*_*/*(2*N*)), regardless of the genetic basis of the trait, provided there is no mutation bias or migration from differently adapted populations.

In contrast, the effect of stabilizing selection on trait variance is mediated by the effect sizes of trait loci. Stabilizing selection reduces trait variance in two ways: first, by disfavouring polymorphism at individual trait loci, and second, by disfavouring multi-locus combinations with a higher than average number of alleles with the same (say, positive) effect on trait value. Selection against polymorphism causes a trait locus with effect size *α* to behave as an underdominant locus with selection coefficient 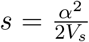 (Robertson, 1956). Selection against multi-locus combinations results in negative fitness epistasis between trait-increasing (or trait-decreasing) alleles, which generates negative LD between trait loci. This kind of negative LD, in turn, has three consequences. First, it causes the genetic variance (i.e., variance in trait values across individuals) to be lower than the genic variance, both within demes and across the full population. Second, it effectively weakens underdominant selection at each locus, thus slightly increasing heterozygosity at trait loci and the genic variance. Third, it reduces the long-term genetic contribution of immigrants to variation within demes, especially under conditions of low gene flow when different demes support different multi-locus combinations.

Below, we compare simulation results with analytical predictions that either neglect the effects of LD on genic variance (LE predictions), or account for it via effective parameters (LD predictions). As detailed in the Methods, the LE prediction treats each trait locus as evolving independently under underdominant selection, allowing us to use the single-locus analysis of the previous section to predict genic variances. The LD predictions then build upon this by assuming that LD due to stabilizing selection essentially alters the effective selection coefficient *s*_eff_ at each trait locus by a factor that depends on the within-deme genic variance 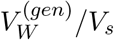, while LD due to migration changes the effective migration rate *m*_eff_ by a factor that depends on the among-deme genic variance 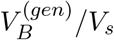.

The main question we address here is: how much more trait variation can be maintained in a structured population compared to an unstructured one? To this end, we always scale trait variances by *V*_*\**_ = 4*µLV*_*s*_, the expected genic variance in a large panmictic population under mutation-selection balance (neglecting LD). In the main text, we keep *Nµ* = 0.01 fixed. In Section SI.f, SI, we consider other values of *Nµ* and show that for a given *Nm*, various quantities depend primarily on the ratio *Nµ/Ns* = *µ/s* (rather than mutation rates and selection coefficients separately).

We first consider traits influenced by *L* loci of *equal* effect size, *α*_*i*_ = *α* for all *i* (and selection coefficient *s*_*i*_ = *s* = *α*^2^*/*(2*V*_*s*_)). Figure 2 shows how the genic and genetic variances change with *Nm*, the average number of migrants per deme, across the full population (Figs. 2a,2d) and within demes (Figs. 2b,2e). Figures 2c,2f show the effect of increasing migration on the expected *F*_*ST*_ at trait loci and the expected *Q*_*ST*_, both of which quantify how genetic variation (at the locus level and trait-level, respectively) is partitioned within and among demes. Symbols show results of individual-based simulations; dashed and solid lines depict LE and LD predictions respectively. The different colours correspond to different strengths of stabilizing selection, *V*_*s*_. However, since *V*_*s*_ only affects evolutionary dynamics via *s* = *α*^2^*/*(2*V*_*s*_), the colours can also be thought of as showing results for traits influenced by loci of large vs. small effect.

#### Genic variances and *F*_*ST*_ at trait loci

First consider how population structure affects 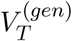, the total genic variance. Figure 2a shows that 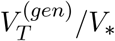 is independent of migration for *Nm < Nm*_crit_, where *Nm*_crit_ observed in simulations is accurately predicted by our LD approximation (and is higher than the critical threshold for a single underdominant locus with 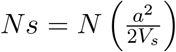; dashed lines). For *Nm < Nm* _crit_, the mean allele frequency in the population at any trait locus is 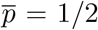, and 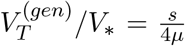 (which is typically much larger than 1), so that the total genic variance in the subdivided population is much higher than that in a large panmictic population. In this parameter regime, different demes occupy different adaptive peaks, so that high levels of variation are maintained in the population as a whole. However, once *Nm* exceeds *Nm*_crit_, the same allele tends to become common across all demes at any trait locus, causing 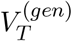 to fall. For large enough migration rates, the subdivided population essentially behaves as a single panmictic population, so that 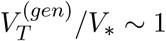.

Figure 2a also shows that population structure increases 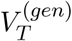 most strongly if *Ns* per locus (and hence, *s/µ*) is large; however, this effect is then restricted to very low migration rates. In contrast, for small *Ns*, the effect of population structure on trait variance is much weaker, but extends over a wider range of migration rates. Thus, the increase in trait variance in populations with (say) *Nm* ≳ 1*/*2 is expected to be modest and entirely due to loci of small or moderate effect.

Next consider how population structure affects the within-deme genic variance, 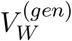. Figure 2b shows that 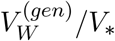 increases with *Nm* at low levels of migration, reaches a maximum at *Nm* ~ *Nm*_crit_, and then begins to decrease as migration increases further, thus mirroring the behavior of within-deme heterozygosity at a single underdominant locus (Fig. 1f). In fact, at *Nm* ~ *Nm*_crit_, the genic variance within a single deme can be as much as five times that in a large panmictic population, provided that *µ/s* ≲ 0.01. More generally, the fact that within-deme genic variance is maximised at *intermediate* levels of migration reflects the tension between two qualitatively different – ‘mixing’ vs. ‘homogenizing’ – effects of migration. For individual demes to maintain high levels of variation, migration must be strong enough to introduce genetic variants from other demes into the focal deme (a ‘mixing’ effect), but not so strong as to drive the same variant close to fixation across all demes (a ‘homogenizing’ effect).

We can also quantify the relative contributions of within-deme and among-deme variation to the overall polymorphism at trait loci using *F*_*ST*_ (Figure 2c). This changes with *Nm* in essentially the same way as the *F*_*ST*_ at a single underdominant locus (see also Fig. 1g). In particular, for any locus with 4*Nm*_crit_ *<* 1 (i.e., *µ/s* ≲ 1*/*24, eq. 22; e.g., green and purple curves in fig. 2c), the average *F*_*ST*_ is higher (lower) than the neutral expectation (black dashed line), depending on whether 4*Nm* is lower (higher) than ~ 1. This observation is interesting in light of previous work that predicts that spatially uniform stabilizing selection should cause trait loci to become more strongly differentiated between diverging populations than neutral loci (Latta, 1998; Le Corre and Kremer, 2012). Our analysis suggests that this is only true if populations evolve largely independently, with even moderate levels of migration being enough to reverse the pattern.

Figures 2a-2c show that while LE predictions (dashed lines) capture the qualitative behaviour of genic variances, predictions that account for LD using effective parameters (solid lines) are more accurate, especially for moderate selection per locus (*Ns* = 1).

Both types of predictions deviate modestly from simulation results for very weak or strong selection. For weak selection (*Ns* ≪ 1), the discrepancy between simulations and theory is a consequence of the relatively small number of demes (*D* = 100) in simulations, which leads to large fluctuations in 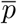, the mean allele frequency at trait loci. As in the case of a single underdominant locus (Fig. S2), the agreement between simulations and theory improves with increasing total population size. For strong selection (*Ns* ≫ 1), theoretical predictions become less accurate for a different reason. If selection per locus is strong and the total mutational input per deme (*NµL*) is low, then very few trait loci are polymorphic within a deme. For instance, for *Nm* = 1, *Ns* = 5 and *NµL* = 2 (in fig. 2), only 6.4% or 1.3% (i.e., ~ 13 or ~ 3) of the *L* = 200 loci are expected to have minor allele frequencies greater than 0.01 or 0.05.Thus, the basic assumptions underlying our theoretical approximations – that trait means hardly deviate from the trait optimum and that trait values are normally distributed – are likely to be violated. Based on this reasoning, we also expect predictions to become more accurate as the number of polymorphic loci increases. To test this, we simulated a single deme under stabilizing selection for different values of *NµL*, while holding constant the scaled parameters *Ns* and *Nµ* (which govern single-locus dynamics) and *µL* (which determines the extent of LD in the *NµL* → ∞ limit). Simulations results indeed approach LD predictions as *NµL* increases (Fig. S4).

#### Genetic variance within demes and across the entire population

We now explore how population structure and selection affect *V*_*W*_ and *V*_*T*_, the genetic variance within demes and across the whole population, as well as *Q*_*ST*_ = *V*_*B*_*/*(*V*_*B*_ + 2*V*_*W*_), which can be thought of as the quantitative genetic analog of *F*_*ST*_ (Spitze, 1993).

Figure 2d shows 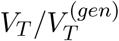, the ratio of the total genetic variance to the total genic variance across the full population, as a function of *Nm* for different strengths of selection. Figure 2e shows the corresponding plot for 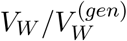, the ratio of within-deme genetic to genic variance. Note that these ratios are always less than 1, which means that, on average, alleles with similar (e.g., positive) effect on trait value are in negative LD with each other.

Negative LD within demes is primarily a consequence of the Bulmer effect, wherein phenotypically extreme individuals, i.e., those with a higher than average number of trait-increasing or trait-decreasing alleles, have lower fitness. This generates negative selection against combinations of such alleles, thus reducing the variance of trait values. However, this is counterbalanced in every generation by the variance released due to random mating and segregation. This so-called segregation variance is rather large (being equal to half the genic variance) when trait loci are unlinked. Thus, within demes, genetic variance is only modestly reduced relative to genic variance: 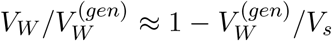 (see also eq. (14)). This also suggests that the effects of LD within demes should be strongest when *Nm* ~ *Nm*_crit_, i.e., when genic variance is at its maximum. This expectation is borne out by Figure 2e, which shows that 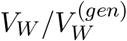 is indeed minimized close to *Nm*_crit_. Further, 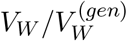 is close to the Bulmer prediction (eq. (14)), especially upon accounting for the effects of LD via effective parameters (solid lines).

In contrast to the variance within demes, genetic variance across the population as a whole can be much lower than the genic variance, i.e., 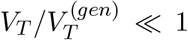, especially at low levels of migration (Fig. 2d). This follows from eq. (21): even when LD within demes is negligible, so that 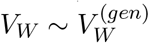, we might still have 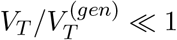, provided *F*_*ST*_ at trait loci is high and *Q*_*ST*_ low. This, in turn, requires that trait means be much less differentiated across demes than the individual loci that contribute to trait variation. This is precisely what occurs at low migration rates – different demes are nearly fixed for alternative alleles at any locus, resulting in high levels of polymorphism across the population as a whole, as well as high *F*_*ST*_ (Figure 2c). However, this does *not* mean that all possible genotypes (that can be constructed by randomly assorting the different alleles) are common in the population. Rather, any deme only supports specific non-random combinations of trait-increasing and trait-decreasing alleles with trait values very close to the trait optimum. This results in strongly negative LD between trait loci at the level of the population as a whole, which constrains trait values and trait means (Figure 2f). Thus, at low migration rates, we have: *Q*_*ST*_ ≪ *F*_*ST*_ ~ 1 and 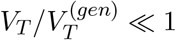.

We also obtain simple predictions for *Q*_*ST*_ (eq. (20)), which match simulations very well (Figure 2f). As expected, *Q*_*ST*_ is markedly lower than the neutral *F*_*ST*_ (dotted black line) – a well-known consequence of spatially uniform stabilizing selection (Leinonen et al., 2013). Another way of seeing this is that *V*_*T*_ ~ *V*_*W*_ (fig. S6), meaning that the total genetic variance across the full population is largely due to variance within demes. Interestingly, while higher values of *Ns* lead to lower *Q*_*ST*_ under very weak migration (*Nm* ≪ 1), the effect of selection becomes weaker with increasing migration. This follows from the fact that for *Nm* ≳ 1, the variance within demes is 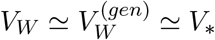, so that:

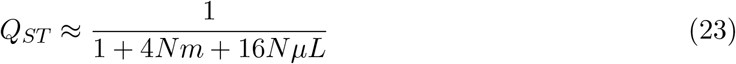

regardless of the strength of stabilizing selection. Thus, *Q*_*ST*_ is significantly reduced relative to neutral *F*_*ST*_ only if 2*NµL*, the average number of new trait-affecting mutations per deme per generation, is comparable to *Nm*, the number of migrants per deme per generation.

### Variance contributions of trait loci with different effect sizes

So far, we have considered traits influenced by loci of equal effect. To evaluate the robustness of our approximations, we now investigate alternative trait architectures. First, consider a simple example where 60%, 30% and 10% of trait loci have weak 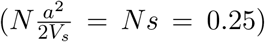, moderate (*Ns* = 1), and large (*Ns* = 4) effects respectively. Figures 3a and 3b show the proportion of genic variance – across the full population and within demes – contributed by each of the three types of loci, as a function of *Nm*.

Under very weak migration, 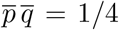 at any locus, regardless of its selection coefficient. Thus, the contribution of any locus *i* to total genic variance 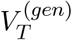 scales with *Ns*_*i*_, meaning that loci of large effect (*Ns* = 4; purple) make disproportionately large contributions to 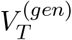 at low migration rates. With increasing migration, the contribution of a given type of locus drops sharply at the associated *Nm*_crit_ threshold (which is lower for larger *Ns*), leading to a compensatory increase in the *relative* contributions of the other types. As migration increases even further, the relative contribution of each type of locus to 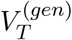 becomes equal to the proportion of loci of that type (indicated by dashed horizontal lines). This is because for very large *Nm*, the structured population behaves as a large panmictic population of size *N*_tot_ = *ND*, in which the average variance associated with any locus is the same (equal to *v*_*\**_ = 4*µV*_*s*_ under LE) for any locus with sufficiently large scaled effect size (i.e., *N*_tot_*s* ~ 1).

The effects of migration on the contributions of different loci to the within-deme genic variance 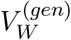 are broadly similar (Fig. 3b). The only difference is that large-effect loci do not contribute as disproportionately to 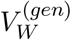 (as they do to 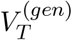) at low migration rates. This reflects the fact that per-locus contributions to 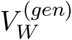 are relatively similar for loci of different effect size (see, e.g., fig. 2b).

We next consider a more realistic trait architecture, where effect sizes are drawn from an exponential distribution with mean (scaled) effect 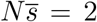 Figures 3c and 3d show the average genic variance contributed by each locus, to the full population and within demes, as a function of its scaled selective effect *Ns*. All variance contributions are scaled by *v*_*∗*_ = 4*µV*_*s*_, the deterministic per-locus variance. As before, we see a good match between simulation results (crosses) and single-locus predictions (dashed lines), with the match improving if we use effective parameters to account for LD (solid lines).

To understand how population structure affects the relative contributions of small vs. large-effect loci to trait variance, first consider the *Nm* = 0 limit (dark blue; Figure 3d). As in the previous example, in this limit, 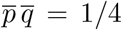 at any trait locus, regardless of its effect size. Thus, loci contribute to the total genic variance (across the full population) in proportion to their effect size (dark blue; Figure 3c). By contrast, the average contribution of a trait locus to the within-deme variance is a non-monotonic function of its effect size: it increases with effect size for low values of *Ns*, peaks at *Ns* ~ 1, and then declines again as *Ns* increases further, reaching its asymptotic value of *v*_*∗*_ = 4*µV*_*s*_ for *Ns* ≫ 1. Thus, with *Nm* = 0, the largest contributions to within-deme trait variance are from loci of intermediate effect (see also Simons et al., 2018). Weak migration between demes (*Nm* = 0.1; light blue) broadens the allele frequency distribution within each deme, thus increasing the contributions of all trait loci to the within-deme variance, regardless of their effect size. At the same time, it depresses the contributions of large-effect loci to the total genic variance. This is because even weak migration is sufficient to drive the same allele to high frequency across the majority of demes at large-effect (strongly selected) loci, thus decreasing the overall level of polymorphism at such loci (fig. 1b).

At even higher levels of migration (*Nm* = 0.5 and 2; pale and dark red), variance contributions of loci with large to moderate effect tend to level out, approaching the deterministic prediction *v*_*∗*_ = 4*µV*_*s*_, while loci of small effect (*Ns* ≲ 0.12 for *Nm* = 0.5, and *Ns* ≲ 0.06 for *Nm* = 2.0) contribute significantly less than *v*_*∗*_. At even larger migration rates, we expect the subdivided population to behave as a panmictic population of size *N*_tot_ = *ND*, in which any locus with *N*_tot_*s* ≳ 1 contributes ~ *v*_*∗*_ to within-deme and total genic variance. In fact, the red curves in Figs. 3c and 3d (corresponding to *Nm* = 2) are already very close to this panmictic expectation (dashed black curves), suggesting that even moderate population structure has little to no effect on the *expected* variance contributions of trait loci.

In addition to measuring per-locus variance contributions, we also track *F*_*ST*_ at trait loci as a function of *Ns* (Figure 3e). As in the case of a single underdominant locus (Fig. 1g), *F*_*ST*_ at trait loci increases with their effect size only in highly structured populations (*Nm* = 0 and 0.1 in fig. 3e). Even modest levels of gene flow (*Nm* ≳ 0.25) are enough to reverse this pattern, causing *F*_*ST*_ to be lower for loci of larger effect.

### Distribution of variances across trait loci

The previous sections focused on the *expected* variance contributions of trait loci. However, the variance *v* contributed by any locus will fluctuate stochastically about this expected value over time and also across demes (in the case of within-deme variance). Thus, we can study the probability *distribution P* (*v*) of *v* or, alternatively, the cumulative probability 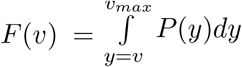, where *v*_*max*_ is the maximum possible variance (corresponding to *p* = 1*/*2). For a locus with selective effect *s* which is polymorphic in a given deme, the within-deme variance *v* can range from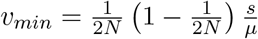 to 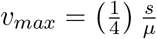(in units of *v*_*∗*_).

We now investigate the behaviour of *F* (*v*), with the aim of characterising ‘outliers’, i.e., loci with unusually large contributions to the within-deme genic variance 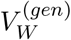. We are interested in outliers because methods for identifying trait loci, e.g., GWAS, are more likely to detect loci with variance above a certain threshold (which decreases with increasing study size; Simons et al. (2018); Spence et al. (2026)). Figure 4a shows the cumulative distribution *F* (*v*) of variance contributions of all trait loci, for a trait influenced by *L* = 200 equal-effect loci with scaled effect size *Ns*, for three different values of *Ns* (colors) and *Nm* = 2. As before, all variances are scaled by the deterministic expectation, *v*_*∗*_ = 4*µV*_*s*_. Note that *v*_*min*_, the minimum possible variance associated with a polymorphic locus, is higher for loci of larger effect, while *F* (*v*_*min*_), the probability that the locus is polymorphic, is correspondingly smaller. Thus, the curves start at different points along the xand y-axes.

**Figure 4.**
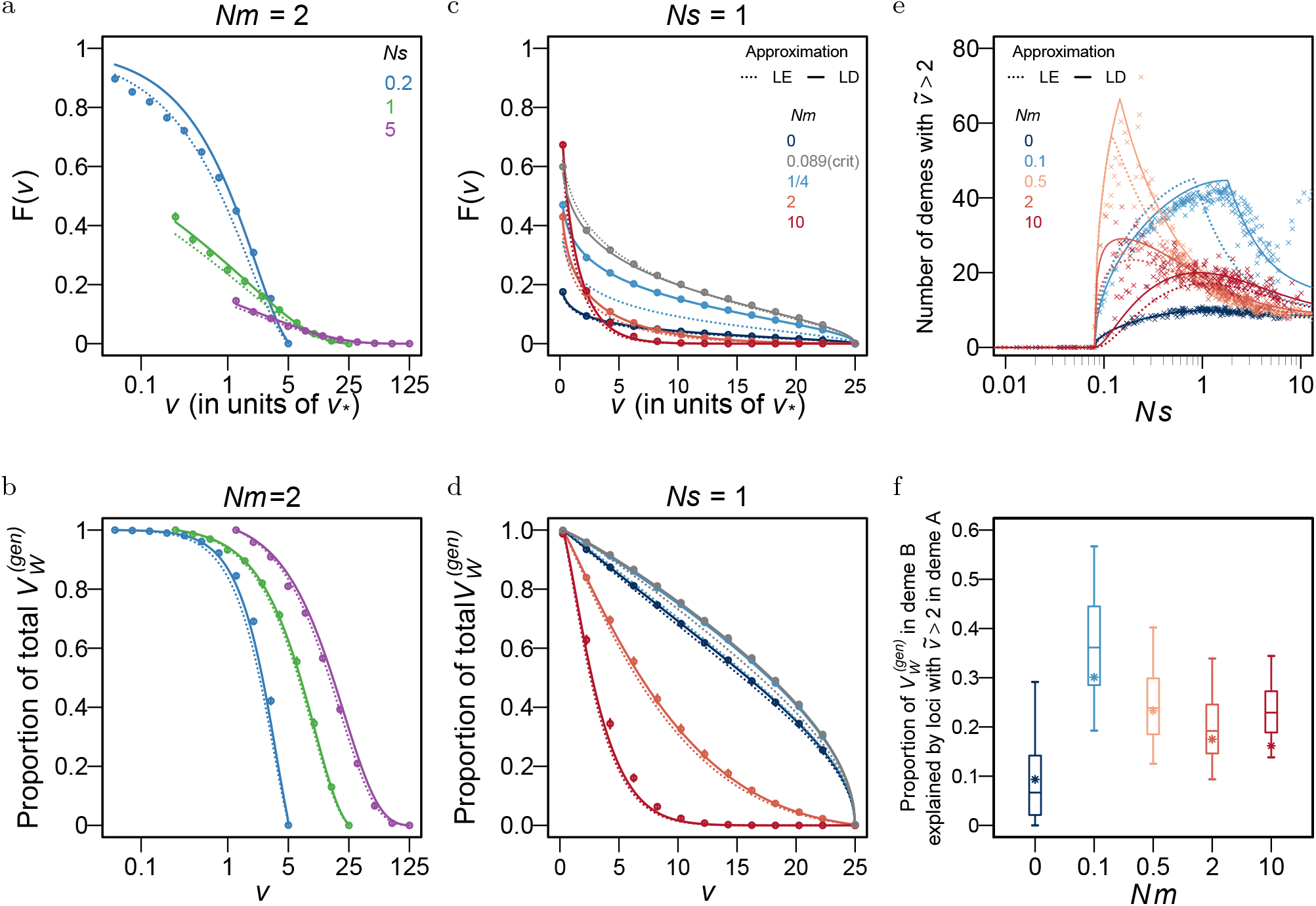
Distribution of the within-deme variance contributions of trait loci. (a),(c) Cumulative probability *F* (*v*) that the variance contributed by a locus exceeds *v*, and (b),(d) proportion of within-deme genic variance 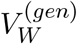 explained by loci with variance exceeding *v*, as a function of *v*, for a trait influenced by *L* = 200 loci of equal effect. Panels (a) and (b) show results for different values of *Ns*, the (scaled) locus effect size, for *Nm* = 2. Panels (c) and (d) show results for different values of *Nm*, for *Ns* = 1. All per-locus variances are measured in units of *v*_*∗*_ = 4*µV*_*s*_, and can range from 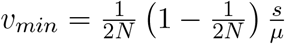 to 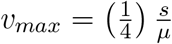 (in these units) for a locus with scaled effect *Ns*. Points show results of individual-based simulations with *N* = 200, *D* = 100 and *Nµ* = 0.01; solid and dashed lines show LE and LD predictions respectively. Panels (e)-(f) show results for a trait influenced by *L* = 200 loci with exponentially distributed effect sizes with mean (scaled) effect *Ns* = 2 (same distribution and simulations as in Figs. 3(c)-(e)). (e) Expected number of demes in which a locus of effect size *Ns* makes a variance contribution exceeding 2*v*_*∗*_, as a function of *Ns*, for different values of *Nm*. (f) Proportion of genic variance in deme B explained by loci with variance contributions greater than 2*v*_*∗*_ in deme A, for different values of *Nm*. Box plots show the 0.05, 0.25, 0.5, 0.75 and 0.95 quantiles of the distribution of the explained proportion of genic variance across different demes in individual-based simulations; the asterisks mark the theoretical LD prediction for the expected proportion.

A striking feature of this figure is that the distributions differ markedly across traits influenced by loci of small vs. large effect size, even though the mean of the distribution (the expected variance) is very similar, being 1.24*v*_*∗*_, 1.19*v*_*∗*_ and 1.09*v*_*∗*_ for loci with *Ns* = 0.2, 1 and 5 respectively in fig. 4a. Broadly speaking, larger effect sizes result in a higher probability of more extreme, i.e., either zero or very large (*v/v*_*∗*_ ≫ 1), variance contributions. Conversely, loci with intermediate contributions (say, 1 *< v/v*_*∗*_ *<* 5) are associated with lower *Ns* values. For instance, in fig. 4a, a locus with *Ns* equal to 0.2, 1 or 5 has a probability 0.45, 0.16, or 0.075 (computed as *F* (5) ™ *F* (1)) of making a contribution 1 *< v/v*_*∗*_ *<* 5 to 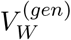.

One can further ask: what proportion of 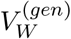 is due to loci with unusually high variance contributions? In other words, when do a small number of such outliers account for most of the variance within demes? Figure 4b shows the proportion of within-deme genic variance explained by loci with contributions exceeding *v* as a function of *v*. Comparing the curves for the various *Ns* values, we see that outliers contribute more when effect sizes are larger (or equivalently, stabilizing selection is stronger). For instance, loci with variance exceeding 5*v*_*∗*_ explain ~ 83% of 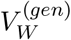 if scaled effect sizes are *Ns* = 5 (which corresponds to *µ/s* = 0.002) and 60% of 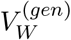 if *Ns* = 1 (i.e., *µ/s* = 0.01).

We also explore how migration between demes affects the contribution of outlier loci to the within-deme trait variance for a trait with intermediate-effect loci (*µ/s* = 0.01; Figs. 4c and 4d). Low levels of migration result in more outliers (as well as a higher proportion of variance explained by such outliers) compared to no migration, provided that *Nm < Nm*_crit_. However, a further increase in migration (beyond *Nm*_crit_) shifts *F* (*v*) towards lower values of *v*, thus decreasing the contribution of outliers to the within-deme variance. Nevertheless, even for migration as high as *Nm* = 10 (dark red curve in Fig. 4c), a significant fraction (~ 17%) of loci have variance contributions exceeding 2*v*_*∗*_ within demes.

Finally, we also investigate the distribution of per-locus contributions to the total genic variance 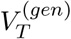 (Fig. S3). As expected, this distribution is quite narrow and concentrated about the expected value 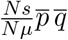 in a population of large total size *N*_*tot*_ = *ND*. More precisely, the width of the distribution depends on the variance of *p*, which approaches zero as *N*_*tot*_ → ∞. What implications do these results have for mapping the genetic basis of trait variation in structured populations? Returning to the example of trait loci with exponentially distributed effect sizes (Figs. 3c-3e), we ask: in how many (out of the 100) demes, does a given locus make an atypically large variance contribution (say, *>* 2*v*_*∗*_)? Figure 4e shows that loci with large contributions to multiple demes tend to be associated with intermediate *Ns* values. If we chose an even larger threshold (e.g., 5*v*_*∗*_), then the typical *Ns* values associated with outliers would be higher, and any locus would be an outlier across fewer demes. However, regardless of the exact threshold for designating outliers, the probability that a locus is an outlier (possibly across multiple demes) is quite sensitive to *Ns*, even for *Nm* as large as 2 in fig. 4e. This contrasts with the average variance per locus, which is largely independent of *Ns* for *Ns* ≳ 0.1 and *Nm >* 1 (fig. 3d). Thus, even moderate to weak population structure can significantly influence the effect size distribution of detected QTL.

We also ask: what proportion of the genic variance in deme *B* is explained by loci identified as outliers (those with variance contributions *>* 2*v*_*∗*_) in another deme A (Fig. 4f)? Interestingly, outliers in A explain the most variance in B at intermediate levels of migration. This is consistent with Fig. 4d, where we see that the relative contribution of outliers to genic variance is maximised close to the associated *Nm*_crit_ threshold. This means that trait variation is *less* polygenic, i.e., is explained by fewer loci with larger individual contributions, if *Nm* ~ *Nm*_crit_. This causes the genetic basis of variation to also become more similar across different demes.

## Discussion

This paper investigates how genetic variation is maintained in a polygenic trait under stabilizing selection in a subdivided population. We show that the diffusion approximation for allele frequencies, coupled with basic quantitative genetic theory, can accurately predict the combined effects of selection and structure on quantitative genetic variation, at least under a simple island model of population structure. Moreover, the effects of LD can be absorbed by effective migration and selection coefficients, allowing for a succinct description.

### How much (more) variation in structured populations

We find that strong population structure can significantly increase diversity at trait loci across the population as a whole by allowing different alleles to persist at high frequency in different demes (fig. 2a). However, since alleles only occur in specific multi-locus combinations (with trait values close to the optimum), trait variance remains quite constrained and is only modestly increased at intermediate levels of migration (fig. S6).Thus, under spatially uniform stabilizing selection, (strong) population structure is mainly a source of cryptic variation that may be ‘unlocked’, e.g., via experimental crosses between individuals from different subpopulations.

These conclusions are in line with those of other models with two demes (Phillips, 1996; Lythgoe, 1997; McDonald and Yeaman, 2018) or of continuous space (Goldstein and Holsinger, 1992; Lande, 1991). In addition, our analysis furnishes an analytical prediction for the critical migration threshold, *Nm*_crit_, below which trait loci exhibit bistability and high levels of polymorphism. Under the island model, we find: 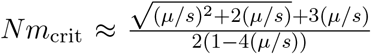 for *Ns* ≲ 1 and 2(*µ/s*) *<* 1*/*2, where 2(*µ/s*) is also the expected heterozygosity at a trait locus in a panmictic population. Thus, in a scenario where typical heterozygosity per trait locus is (say) ≲ 0.1 in a panmictic population, *Nm* between demes must be at most ~ 0.3 for population structure to significantly increase trait variation. Thus, broadly speaking, the rather weak levels of population structure found in most natural populations (see, e.g., Morjan and Rieseberg, 2004) cannot significantly inflate quantitative genetic variation, at least under spatially uniform stabilizing selection and with island-model migration (i.e., no isolation by distance).

There are two caveats to this. First, we focus on populations at mutation-selection-migration-drift *equilibrium*, which may not be attainable over biologically relevant timescales. For instance, imagine a scenario where a set of long-isolated populations (each with its own set of trait alleles) come into contact. While equilibrium predictions suggest that the same allele should eventually spread across all subpopulations at any locus of sufficiently large effect, this requires populations to cross a ‘fitness valley’ (arising from selection against rare alleles at any locus), which can take very long under weak migration (see also Fig. S5). Thus, structured populations may maintain much more variation than predicted by equilibrium theory.

Second, even though the *expected* variance contributed by (most) trait loci hardly changes beyond *Nm* ~ 2 (see, e.g., Figs. 2b and 3d), the *distribution* of variance contributions is sensitive to much higher levels of migration (fig. 4c,d). For instance, in fig. 4d, we see that the proportion of 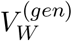 explained by loci with atypically large contributions is very different for *Nm* = 2 vs. *Nm* = 10. This further supports the idea that spatially restricted sampling of individuals (e.g., from a local subpopulation that is embedded in a larger population) may give a rather different picture of trait architecture compared to more widespread sampling (see also Battey et al., 2020; Steiner et al., 2025), even without local adaptation and under rather weak population structure.

### How much differentiation between subpopulations

We analyse differentiation between subpopulations in two ways. We first ask whether the same (or alternative) alleles occur at high frequency in different demes. We then consider *F*_*ST*_ and *Q*_*ST*_, which quantify the relative contribution of among-population differences to genic diversity (at trait loci) and trait variance respectively.

In general, we expect different sets of alleles to be nearly fixed in populations with a long history of complete isolation (*Nm* ~ 0). This process of allelic differentiation between populations is actually accelerated by stabilizing selection, which increases the rate at which minor alleles are lost (Yair and Coop, 2022). As discussed above, migration between populations alters this expectation, causing the same allele to reach high frequency across all demes, with the critical migration threshold, *Nm*_*crit*_, at which this occurs being lower for loci of larger effect. A second observation is that while *F*_*ST*_ at trait loci is always higher than the neutral expectation under weak migration, this pattern is actually reversed at higher levels of migration (Fig. 3e).

These findings bear upon recent studies that aim to distinguish between alternative modes of selection (purifying vs. stabilizing vs. none) based on expected allele sharing at putative QTL between humans of different ancestries (Patel et al., 2024). These studies neglect gene flow between ancestries, which is known to have occurred (see, for instance Jouganous et al., 2017). This may bias conclusions, since, as we show here, even moderate levels of gene flow can affect allele sharing (e.g., as measured by *F*_*ST*_) at selected loci (see, e.g., fig. 3e). Moreover, since sites assayed in GWAS are often not causal themselves (but only in imperfect LD with the causal locus), gene flow may also affect the strength of LD between the assayed site and the causal locus it tags, thus further complicating interpretation.

Our results are also relevant to recent work exploring whether introgression between populations adapted to the same trait optimum may be impeded by stabilizing selection in the recipient population (Veller and Simons, 2024; Ragsdale, 2025). Such selection disfavours descendants of migrant individuals, that are *more* phenotypically variable than residents, provided 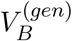 is high (see also eq. (16)). However, as we show here, even low levels of migration are enough to drastically reduce among-population differentiation. Thus, spatially uniform stabilizing selection is unlikely to generate a significant barrier to gene flow unless populations have a long history of nearly complete isolation.

### Do the same or different alleles underlie trait variation in different subpopulations

We find that an allele is more likely to be significantly associated with trait variation across different subpopulations when migration between populations is moderate (Figs. 4e, 4f). This is because very low or very high migration between demes drives alleles towards local or global fixation respectively. This shifts the distribution of per-locus variance contributions to lower values, making it less likely that a locus will contribute significantly to multiple demes. This contradicts the general expectation that higher migration between populations should necessarily lead to more shared standing variation, thereby increasing parallel adaptation, e.g., in response to changing environments. Here, we demonstrate that while high migration necessarily increases allele sharing (by causing the same alleles to nearly fix) across sub-populations, it actually decreases shared polymorphism and standing variation (4f).

What does this mean for mapping the genetic basis of variation in extended populations? Our results suggest that spatially limited sampling of individuals may actually be more useful (than broad sampling) for identifying trait loci. This is because the typical variance contributions of trait loci tend to be higher within individual subpopulations than across the population as a whole, especially for loci of large effect (contrast, e.g., Figs. 4a,b with fig. S3). This phenomenon is well known in the context of GWAS, where alleles significantly associated with trait variation in one population are usually not globally significant. This limits the ‘portability’ of so-called polygenic scores (and other methods of genomic prediction) across populations (Duncan et al., 2019; Wang et al., 2020). As found by Yair and Coop, 2022, stabilizing selection further reduces portability by reducing shared standing variation. However, here we show that this effect is quite sensitive to migration between populations (fig. 4f), with intermediate levels of gene flow significantly mitigating the effects of stabilizing selection on shared variation.

### Towards a more quantitative interpretation of Q_**ST**_ **vs. F**_**ST**_

In agreement with previous work (reviewed in Leinonen et al., 2013), we find that spatially uniform stabilizing selection reduces variation in trait means across subpopulations, causing *Q*_*ST*_ to be lower than neutral *F*_*ST*_ (fig. 2f). In addition, we obtain a simple analytical prediction under the island model: 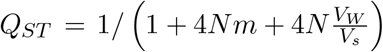(see eq. (20)). Here, *V*_*W*_ */V*_*s*_ approaches 4*µL* (which is twice the total mutation rate for the trait) with increasing migration. This suggests that the difference between *Q*_*ST*_ and neutral *F*_*ST*_ should be pronounced when 4*µL* ≫ *m*, i.e., if population structure is strong and/or selected traits have large mutation targets. Moreover, this difference should be largely independent of both the strength of stabilizing selection and the effect sizes of trait loci (provided that *Nm* is not too low).

Thus, in theory, comparing 1*/Q*_*ST*_ with 1*/F*_*ST*_ (at neutral loci) can furnish an estimate of 4*NµL*, the total mutational input for the selected trait, or at least, provide a means of comparing mutation rates across traits. However, obtaining reliable estimates of *Q*_*ST*_ is often difficult in practice (Whitlock, 2008). Also, genetic correlations between traits would likely alter our simple prediction for *Q*_*ST*_ for the focal trait, which will now be influenced by (spatially uniform or divergent) selection on other correlated traits. Thus, extending the model to multiple traits is a fruitful direction for future work that could more broadly clarify the extent to which *Q*_*ST*_-*F*_*ST*_ comparisons can provide information about (multivariate) selection in structured populations. As noted by Le Corre and Kremer (2003), comparing *Q*_*ST*_ to the average *F*_*ST*_ across quantitative trait loci (which they refer to as *F*_*ST Q*_) also gives information about the proportion of trait variance that is due to LD between trait loci (see also our eq. (21)). In particular, we find that even when LD within demes is negligible, there can be substantial negative LD in the population as a whole, provided that *F*_*ST*_ at trait loci is high and *Q*_*ST*_ low, which, in turn, requires *Nm* ≲ 1 (fig. 2d). Thus, while stabilizing selection only contributes weakly to LD in panmictic populations (unless trait loci are tightly linked), the combination of stabilizing selection and strong population structure can generate strong negative LD, causing genetic variance across the whole population to be up to an order of magnitude lower than genic variance (fig. 2d).

### Extensions and open questions

We consider a simple model of selection in which fitness depends on a single polygenic trait under spatially uniform stabilizing selection. However, in reality, genetic variants may affect multiple traits, some of which may be subject to spatially uniform and others to divergent selection. Such a scenario, while highly relevant to evolutionary inference, remains under-explored in theoretical work (though see Guillaume and Whitlock (2007); MacPherson et al. (2015)). In particular, we lack analytical predictions for how selection on multiple correlated traits and gene flow between populations together shape genetic variances and covariances across traits and loci. The diffusion approximation can be readily extended to allow for a more general relationship between the selection coefficient at a given locus and its effect on a focal trait (see, e.g., Simons et al. (2018)). Thus, the theory developed here provides a natural starting point for investigating migration-selection equilibrium across many traits.

We also make various simplifying assumptions about trait architecture, namely that trait loci are unlinked, have additive effects (no phenotypic dominance) and that there is no mutation bias (so that expected trait means coincide with the trait optimum). The last assumption is crucial, since biased mutation (say, towards trait-increasing alleles) would cause the trait mean to deviate from the trait optimum, thus generating directional selection (in addition to underdominance) at each locus. Both directional selection and mutation bias can increase the frequency of alleles with, say, positive (compared to negative) effect on trait value (Charlesworth, 2013): thus, it is unclear if their signatures can be distinguished in genomic data (Koch et al., 2024; Berg et al., 2025). Our analytical mean-field approach based on the diffusion approximation can potentially be extended to explore this question.

Finally, we consider a very simple island model of population structure, which has the advantage of being analytically tractable, thus allowing for detailed prediction. Such many-deme models represent at least as (un-)realistic a reference point for natural populations as two-deme models, which underlie the bulk of theory on selection in structured populations. However, both many and two-deme models fail to capture the complexity of selection across continuous space, wherein allele frequencies change over a characteristic spatial scale determined by the balance between dispersal and selection per locus (Barton, 1999). For weakly selected loci, this spatial scale may be comparable to that associated with neutral isolation-by-distance (Nagylaki, 1978), making it difficult to distinguish selected loci from the neutral background. Understanding to what extent signatures of (spatially uniform vs. heterogeneous) selection can be disentangled from those of spatial population structure in genomic data thus remains an open question, in sore need of more theoretical work.

## Data Availability

Mathematica notebook with all numerical computations and Python code for individual-based simulation are available at https://github.com/lijuan2010big/PopulationStructureAndStablizingSelection-Equilibrium.git.

## Acknowledgments

We thank Nick Barton, Vitor Sudbrack and colleagues of our joint project (SFB polygenic adaptation) for helpful discussion and comments on the manuscript. This work was supported by part of SFB Polygenic adaptation (FWF 10.55776/F91) to HS. The computational results presented have been achieved in part using the Austrian Scientific Computing (ASC) infrastructure.

## Supplementary information

### SI.a Weak and strong selection approximations for a single underdominant locus

In principle, the equilibrium allele frequency distribution in eq. (7) fully specifies the behaviour of a single underdominant locus at mutation-selection-migration-drift equilibrium under the island model. However, this distribution 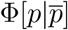 is a function of 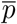, the mean allele frequency across the whole population, which is itself obtained by numerically integrating over 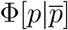, thus necessitating a numerical solution. In this Appendix, we derive simpler analytical expressions for various quantities by considering the weak selection and strong selection approximations of the full equilibrium distribution.

For clarity, we re-write equation (7) below, omitting the subscript *i* for the locus (since we are only considering a single locus) as well as the subscript *j* for the deme. Then we have:

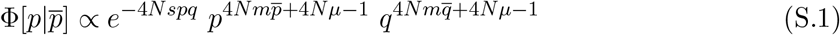

where 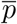 can be found by numerically solving:

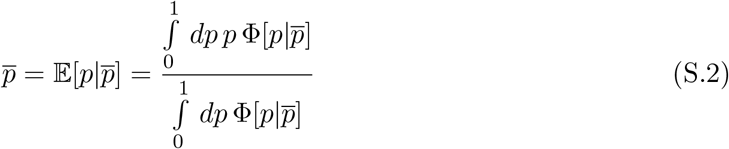

#### Weak selection

For *Ns* ≪ 1, we can approximate the allele frequency distribution (eq. (S.1)) as: 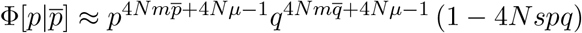. Substituting into eq. (S.2), and retaining only first-order terms in small parameters (i.e., in *Ns* and *Nµ*) yields a cubic equation for 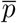. This has the following three solutions:

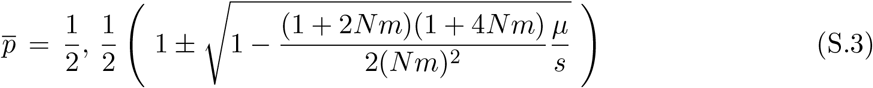

By evaluating the stability of the three equilibria, we find that there is a critical migration threshold *Nm*_crit_, such that 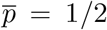 is the only stable equilibrium for *Nm < Nm*_crit_ and 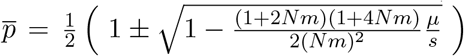 are the only stable equilibria for *Nm > Nm*_crit_. The critical migration threshold (at which the stability of the equilibria changes) can be obtained as the *Nm* value for which the expression under the square root in eq. (S.3) becomes zero. This yields:

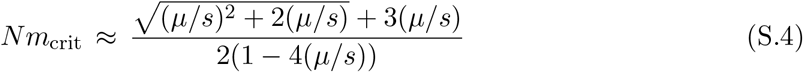

Note that for *µ/s* ≪ 1, eq. (S.4) reduces to 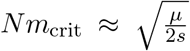, which is eq. 25b of Barton and Rouhani (1993). However, eq. (S.4) is more accurate for larger values of *µ/s* (in practice, for *µ/s >* 0.01), and applies as long as *µ/s <* 1*/*4 (which is the deterministic criterion for bistable allele frequencies at an underdominant locus in a panmictic population).

Equation (S.3) can also be used to calculate the critical selection strength *Ns*_crit_ (above which bistability occurs) as a function of *Nm*, at least in a parameter regime in which *Ns*_crit_ ≪ 1 (which requires *Nm* to be reasonably large). This is simply given by:

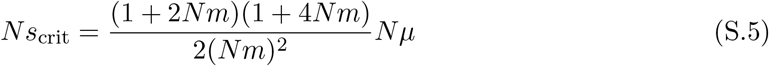

The above threshold diverges as *Nm* → 0 (and is thus not accurate in the low-migration regime). At the other extreme of high migration, we have *Ns*_crit_ ≈ 4*Nµ*, or *s*_crit_ = 4*µ*, which is the critical selection threshold for bistability in a large panmictic population.

We can also calculate 𝔼[*pq*] by explicitly integrating over the weak selection approximation for Φ[*p*]. Then, using the expression for 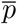 in eq. (S.3), and retaining terms up to first order in *Ns* and *Nµ*, gives the following expression for 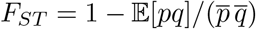:

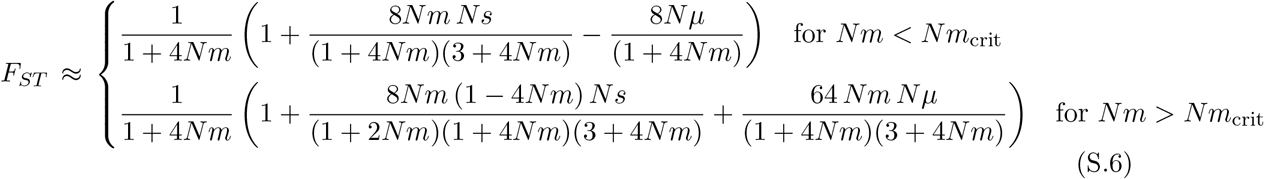

Equation (S.6) shows that for *Nm > Nm*_crit_ and in the limit *Nµ* → 0, the first-order correction to *F*_*ST*_ due to selection changes sign precisely at *Nm* = 1*/*4. Thus, *F*_*ST*_ is greater than 1*/*(1 + 4*Nm*) (the neutral expectation) for *Nm <* 1*/*4, and less than 1*/*(1 + 4*Nm*) for *Nm >* 1*/*4, provided that *Nm*_crit_ *<* 1*/*4.

#### Strong selection

If selection is strong and migration sufficiently weak, then the equilibrium distribution is sharply concentrated near the boundaries *p* = 0 and *p* = 1. Then we can approximate the distribution near each boundary by a Gamma distribution, such that: 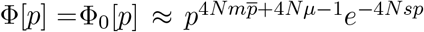 near *p* = 0, and 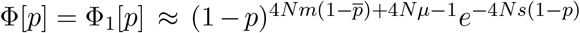near *p* = 1. We then use the fact that Φ_0_(*p*) and Φ_1_(*p*) fall off rapidly to zero on moving away from the boundaries *p* = 0 and *p* = 1 respectively. This allows us to approximate the expected allele frequency as:

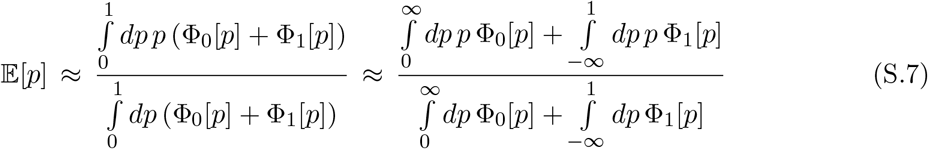

Similarly, the expected heterozygosity can be approximated as:

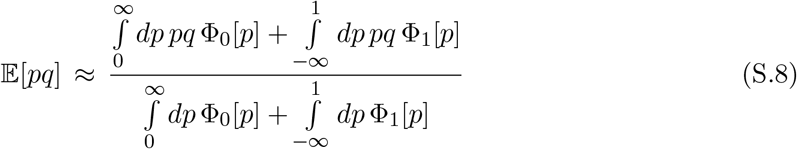

Now using 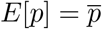 gives the following equation for 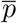:

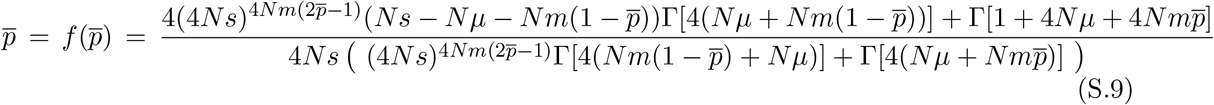

where Γ[*x*] denotes the Gamma function. Equation (S.9) can be solved numerically to obtain the equilibrium allele frequency 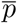. As in the case of weak selection, multiple equilibria are possible, whose stability changes with increasing *Nm*. For *Nm < Nm*_crit_, we have a single stable equilibrium at 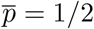, whereas for *Nm > Nm*_crit_, there are two stable equilibria – one lower than and one greater than 1*/*2. One can obtain the critical threshold *Nm*_crit_ by noting that it is that value of *Nm* at which *f* ^*′*^(1*/*2) = 1. This gives:

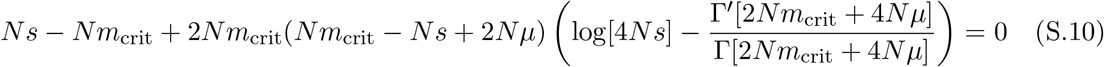

where Γ^*′*^[*x*] denotes the derivative of the Gamma function with respect to *x*. Equation (S.10) can be solved numerically to obtain the critical migration threshold *Nm*_crit_. Alternatively, we can obtain an explicit closed-form expression in the limit where *Nµ* ≪ 1 (which is a natural assumption) and *Nm*_crit_ is also sufficiently small (which requires *Ns* to be large). Then we have: 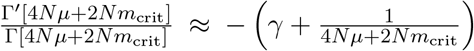. Substituting this into eq. (S.10) and retaining terms up to first order in the small parameters *Nm*_crit_ and *Nµ* gives a quadratic equation for *Nm*_crit_, which can be solved to obtain *Nm*_crit_ as a function of *Ns* and *Nµ*. To lowest order in *Nµ*, this is:

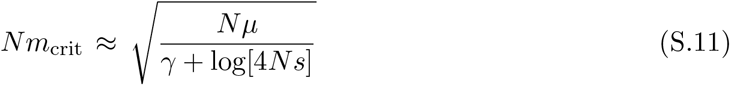

This is the same as eq. 21b of Barton and Rouhani (1993) with an erroneous factor of 2 corrected.

**Figure S1.**
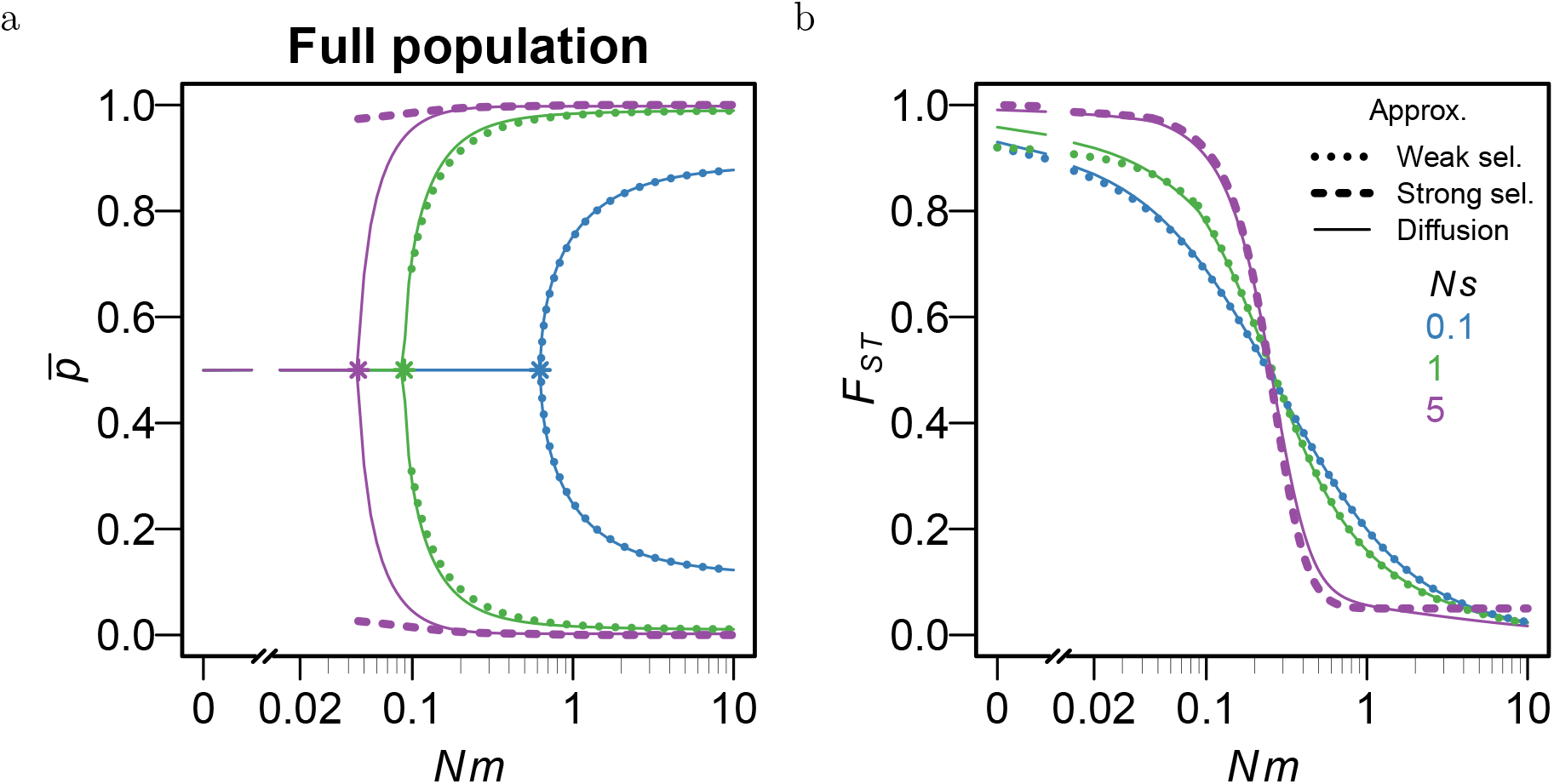
Comparison of the full diffusion approximation (solid lines) with the strong selection (dashed line) and weak selection (dotted line) approximations for a single underdominant locus. (a) Mean allele frequency across the full population 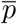 and (b) *F*_*ST*_ at the selected locus as a function of *Nm* for three different values of *Ns*, assuming *Nµ* = 0.01. Asterisks in (a) mark the critical migration rates (*Nm*_crit_) as predicted by the diffusion approximation. Weak selection approximations are reasonably accurate for *Ns* as large as 1.

We can also obtain explicit expression for 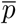 and *F*_*ST*_ when *Nm* is sufficiently larger than *Nm*_crit_ that 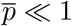, but small enough that the allele frequency distribution can be approximated by the sum of Gamma distributions (as above). Then assuming that 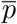 is 𝒪(*µ/s*), Taylor expanding equation (S.9) in powers of *µ/s* and solving the resultant equation gives:

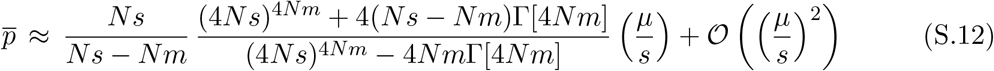

Similarly, we have:

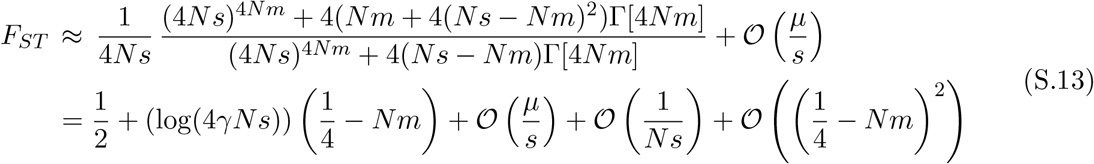

As in the case of weak selection, we see that *F*_*ST*_ exceeds the neutral expectation 1*/*(1+4*Nm*) for *Nm* ≲ 1*/*4 but falls below it for *Nm* ≳ 1*/*4. This change in *F*_*ST*_ occurs at precisely *Nm* = 1*/*4 in the limit of very strong selection, i.e., as *µ/s* → 0 and *Ns* → ∞.

We now test the weak selection and strong selection approximations by comparing *p* and *F*_*ST*_ as obtained numerically from equations (S.1) and (S.2) (i.e., the full diffusion approximation) against the predictions of equations (S.3) and (S.6) (for weak selection) and equations (S.12) and (S.13) (for strong selection). As evident from Figure S1, the weak selection predictions are very accurate for *Ns* = 0.1, and are reasonable even for *Ns* = 1, regardless of *Nm*. By contrast, the strong selection predictions for *p* are only accurate for *Nm* ≲ 1: this is not surprising since the strong selection approximation assumes that allele frequency distribution are concentrated near *p* = 0 and *p* = 1 (which requires migration to be weak).

### SI.b Effective migration rate at a trait locus

In this section, we derive an expression for the effective migration rate *m*_eff_ at a site that is unlinked to any trait locus. This derivation follows the basic argument in Surendranadh and Sachdeva (2025), which we recapitulate here for completeness.

In general, *m*_eff_ *< m* if immigrants and/or their descendants have lower fitness, on average, than the residents in any given deme (where ‘residents’ will be defined more precisely below). In this case, immigrant alleles are selected against, even if they are neutral, due to their association with less fit genetic backgrounds. However, as long as the focal allele is unlinked to any selected locus, such associations break down over ≲ 10 generations – much faster than the timescales that govern single-locus dynamics (which depend on 1*/m*, 1*/s, N*, etc.). Thus, we can describe the effect of indirect selection arising from these transient associations simply by substituting an effective migration rate (instead of the raw migration rate) into the single-locus equations for allele frequency dynamics (Sachdeva, 2022).

In the limit *m* → 0, the ratio *m*_eff_*/m* is simply equal to the average reproductive value of migrants (Kobayashi et al., 2008), which is their long-term contribution to the gene pool of the recipient population, relative to that of residents. Alternatively, *m*_eff_*/m* can be thought of as the factor by which the fixation probability of a neutral migrant allele is reduced relative to that of a neutral resident allele. Further progress can be made by noting that under random mating, the probability of mating between individuals with *recent* migrant ancestry is O(*m*^2^), and thus, can be neglected for *m* ≪ 1. Thus, in this limit, migrant individuals mate primarily with residents to produce *F* 1 hybrids, which mate primarily with residents to produce first-generation backcrosses and so on.

In the following, we first derive an expression for the effective rate of migration from a specific donor deme to a specific recipient deme; we will then take the expectation over all pairs of donor and recipient demes. We will use the subscript ‘0’ to denote a migrant, ‘1’ to denote the first-generation descendant of the migrant (an F1 individual), and *i* = 2, 3, … to denote the *i*^*th*^ generation descendant of the migrant (an (*i* − 1)^*th*^ generation backcross). Further, we will designate any individual within the recipient deme with no migrant ancestor in the last *g* generations as a ‘resident’ (represented by the subscript ‘r’). The cutoff *g* is arbitrary and will later be assumed to approach infinity. Let 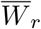and 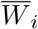denote the mean fitness among residents and among *i*^*th*^ generation descendants of migrants. Then, following Westram et al. (2022), we can express the effective migration rate at an unlinked locus as:

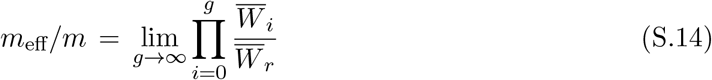

To find the various 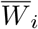, we further assume that trait values within any population sub-group, say the *i*^*th*^ generation descendants of migrants, follow a distribution *P*_*i*_(*z*), which is approximately normal with mean 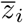 and variance *V*_*i*_. Similarly, resident trait values are also approximately normally distributed with mean *z*_*r*_ and variance *V*_*r*_. We can then express *W*_*r*_ and *W*_*i*_ as:

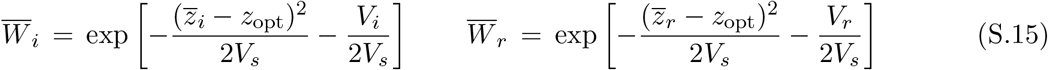

In the next step, we establish how the mean and variance of trait values within any population subgroup are related to those in the parental subgroups. More specifically, one can express the trait value distribution *P*_*i*_(*z*) in the *i*^*th*^ subgroup as:

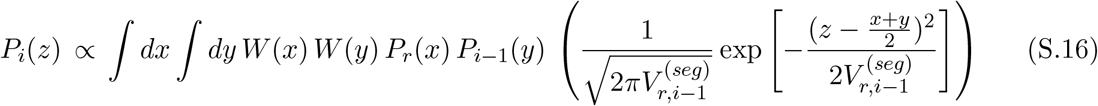

Here, *x* and *y* are respectively the trait values associated with the resident parent and the parent who is an (*i* − 1)^*th*^ generation descendant of a migrant; *P*_*r*_(*x*) and *P*_*i−*1_(*y*) are the corresponding trait value distributions. Parental trait value distributions are weighted by the fitness function 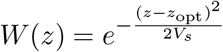. Finally, the last term describes the inheritance of a quantitative trait under the infinitesimal model: the trait value *z* of the offspring is assumed to be normally distributed around the average of the two parents with variance equal to the segregation variance 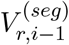 released by mating between residents and (*i* − 1)^*th*^ generation descendants of migrants.

Assuming, as before, that *P*_*r*_(*x*) and the various *P*_*i*_(*y*) are normal distributions, and integrating over *x* and *y* yields the following recursions:

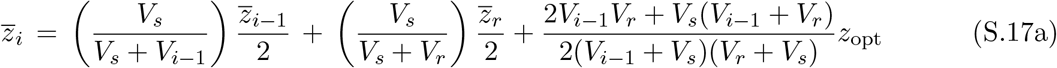

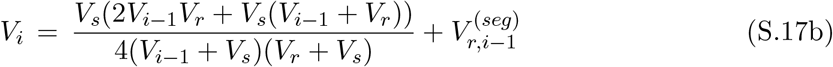

The above equations simplify considerably if *V*_*r*_*/V*_*s*_ ≪ 1 and *V*_*i*_*/V*_*s*_ ≪ 1. Then, to zeroth order in *V*_*r*_*/V*_*s*_ and *V*_*i*_*/V*_*s*_, we have:

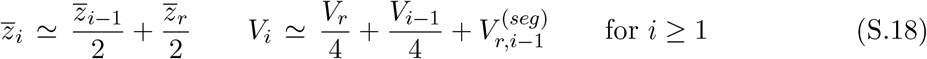

However, even without assuming variances to be small, as long as 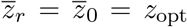, we have: 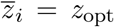, i.e., all population subgroups (regardless of proportion of migrant ancestry) have mean trait value equal to the trait optimum.

In the last step, we establish how the various segregation variances 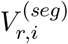 are related to the within-deme and among-deme genic variances. Let *p*_*r,k*_ and *p*_*i,k*_ denote respectively the allele frequency at trait locus *k* among residents and among *i*^*th*^ generation descendants of migrants. As above, if *V*_*r*_*/V*_*s*_ ≪ 1 and *V*_*i*_*/V*_*s*_ ≪ 1, then to leading order in these small terms, we have:

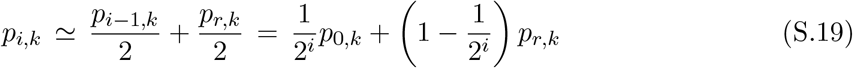

This is simply the change in allele frequencies due to segregation at each trait locus and neglecting the effects of selection.

We can now express the various segregation variances in terms of allele frequencies as:

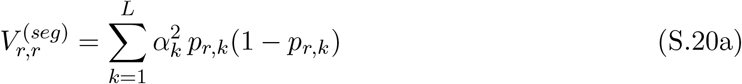

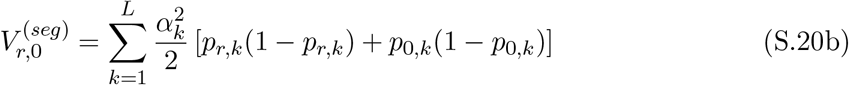

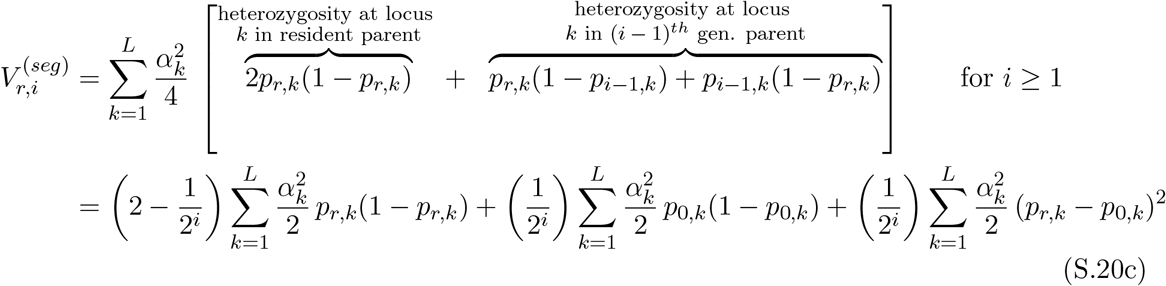

In expectation, we have: 𝔼 [*p*_*r,k*_(1 − *p*_*r,k*_)] = 𝔼 [*p*_0,*k*_(1 − *p*_0,*k*_)] and 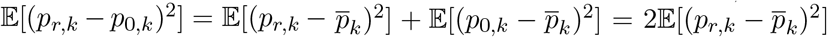 (for the infinite-island model), where *p*_*k*_is the mean allele frequency at locus *k* across the entire population. Thus, in expectation, we have:

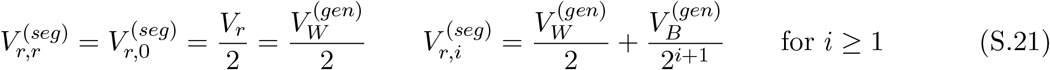

where 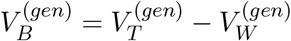 is the among-deme genic variance.

Combining equations (S_.14) and (S.15) with the observation that 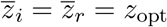 for all *i* (in expectation), we obtain:

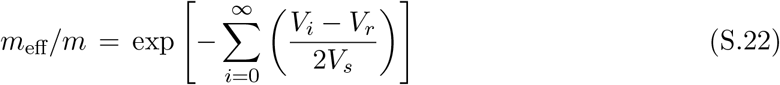

Combining equations (S.18) and (S.21) with the observation that *V*_*r*_ = *V*_0_ gives:

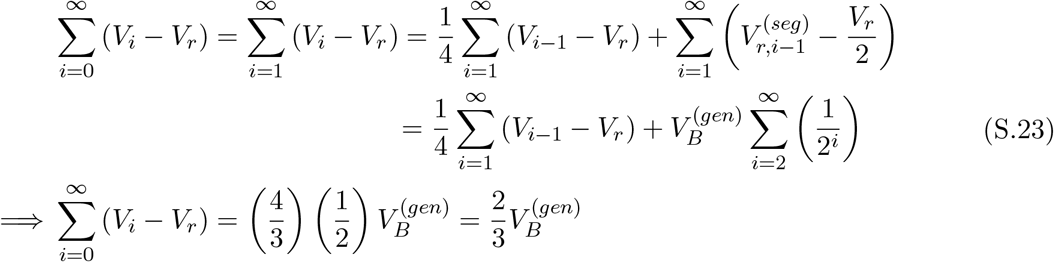

Thus, we finally have:

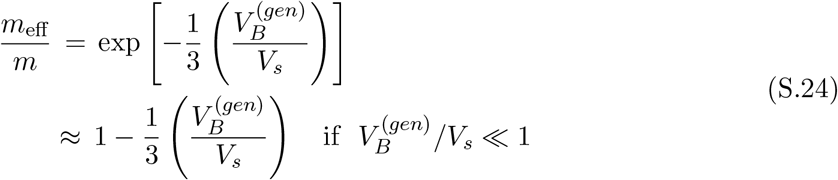

This is the same as equation 16 of Veller and Simons (2024) (which is expressed in terms of *F*_*ST*_ at trait loci instead of 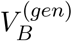 up to a factor of 2. The factor of 2 difference arises from the different (two-island vs. infinite-island) models analysed in the two studies.

The derivation above is strictly valid only for a neutral locus that is unlinked to trait loci. By contrast, *m*_eff_ at a trait locus should also depend on *s*, the strength of direct selection at that locus. However, this effect can be neglected if *s* ≪ 1 (Sachdeva, 2022), allowing us to approximate *m*_eff_ at a trait locus via eq. (S.24). The assumption that *s* ≪ 1 also underlies the diffusion approximation and is not overly restrictive as *Ns* can still be substantial.

### SI.c Factors influencing the accuracy of theoretical approximations

Our theoretical approximations are based on the infinite-island (*D* → ∞) assumption and also rely on the standard quantitative genetic assumption of normally distributed trait values (which is justified in the *L* → ∞ limit). Here, we explore to what extent departures from these assumptions might cause discrepancies between simulations and theory.

#### Finite number of demes

Taking the *D* → ∞ limit (i.e., assuming an infinite number of islands) allows us to treat 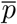, the mean allele frequency at any locus, as deterministic. However, in a finite population, 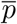 will fluctuate about its expected value, causing the total heterozygosity 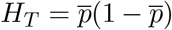 to be lower in expectation than its predicted value (since 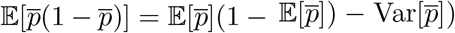). This is indeed what we observe in Figs. 1e for a single underdominant locus under weak selection (*Ns* = 0.1) and for *Nm* less than or close to the predicted *Nm*_crit_. Theory predicts *p* = 1*/*2 for these parameters. However, in simulations, *p* varies considerably around this expected value, especially for *Ns <* 1 and with *D* = 100. This reduces *H*_*T*_ as well as *H*_*W*_, since deviations of 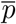 from 1*/*2 also tend to reduce 𝔼 [2*pq*]. A similar pattern is also observed for the polygenic trait (Figs. 2a and 2b) for *Ns* = 0.1.

To investigate this further, we consider a specific parameter combination (*Ns* = 0.1, *Nµ* = and *Nm* = 0.5) for which there is a significant discrepancy between simulations and theoretical predictions for a single underdominant locus in Figs. 1e and 1f. Varying the number of demes *D* while keeping all other parameters fixed (Fig. S2a), we find that *H*_*T*_ and *H*_*W*_ (as measured in simulations) both increase with *D*, approaching the infinite-island prediction as *D* becomes large. This is consistent with the fact that fluctuations in *p* (about the expected value of 1*/*2) fall with increasing *D*.

We then ask: is the variance of 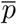 influenced directly by the number of demes or only via its effect on the total population size *N*_*tot*_ = *ND*? To distinguish between these two possibilities, we perform simulations where *D* is increased, while simultaneously decreasing *N*, increasing *m*, and keeping *µ* and *s* fixed (Fig. S2b). Thus, *N*_tot_*µ* = *NDµ* (total mutation rate across the full population), *Nm* (number of migrants per deme) and *µ/s* (strength of mutation relative to selection) remain unchanged as the *D* increases. Figure S2b shows that there is very little effect of *D* on heterozygosities in this case. Further, the discrepancy between simulations and theory is lower for larger values of *N*_tot_*µ* (circles vs. squares). This suggests that for a given value of *Nm* and *µ/s*, the number of demes affects heterozygosities only via *N*_tot_*µ*.

**Figure S2.**
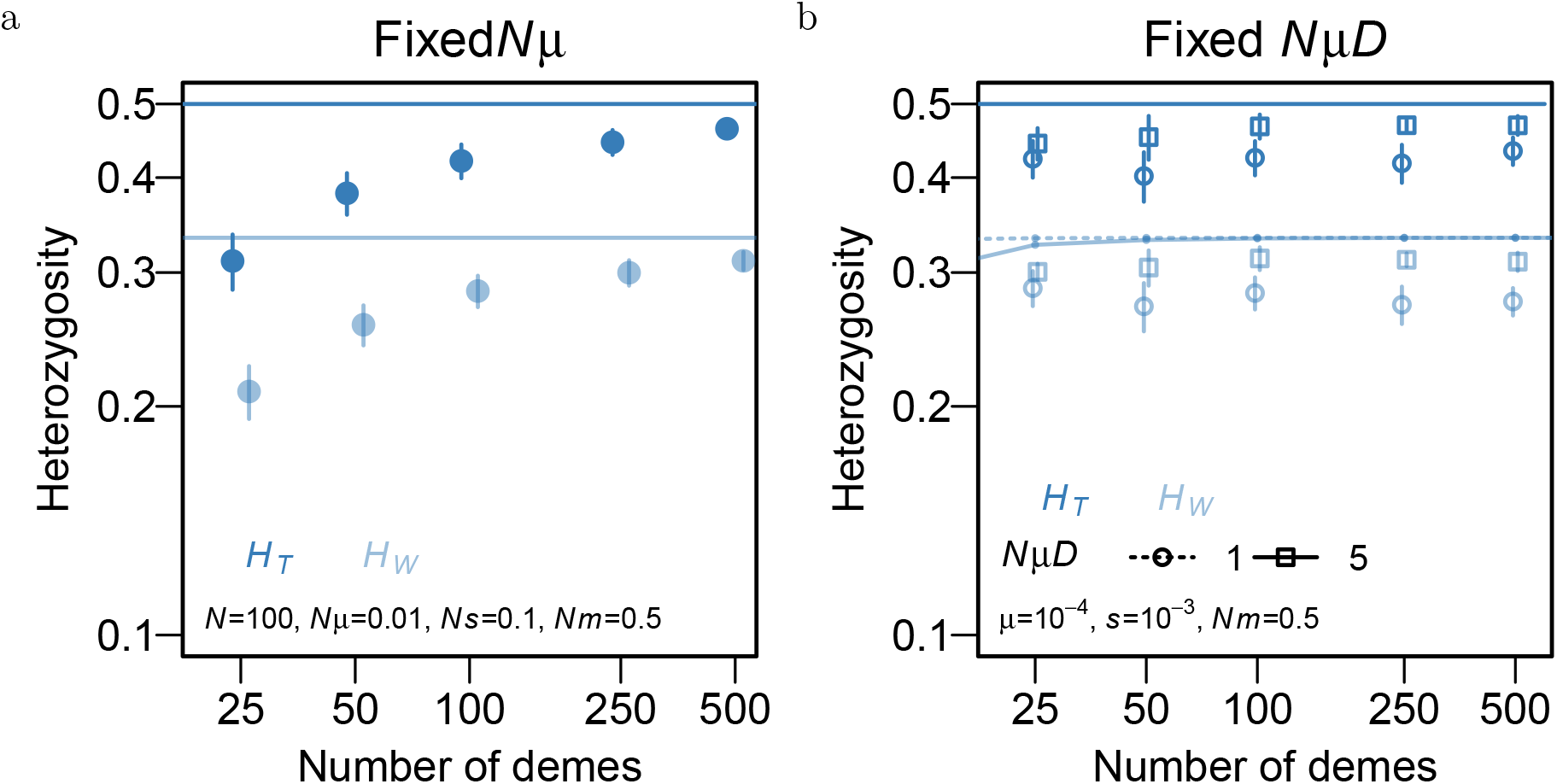
Within-deme heterozygosity (*H*_*W*_; light blue) and total heterozygosity (*H*_*T*_; dark blue) at a single underdominant locus as a function of number of demes *D* with (a) *Nµ* = 0.01 and *N* = 100 held constant (b) *NµD* = 1 (circles) or *NµD* = 5 (squares) constant. Symbols show the results of individual-based simulations while lines indicate theoretical predictions for the infinite-island model. In (b), the mutation rate and selection coefficient are fixed at *µ* = 10^*−*4^ and *s* = 10^*−*3^; the number of demes *D*, the population size per deme *N*, and migration rate *m* are varied simultaneously such that *Nm* and *NµD* remain constant. Simulation results approach the infinite-island prediction with increasing total population size *N*_tot_ = *ND* (in both (a) and (b)).

In addition to tracking the expected *H*_*T*_ and *H*_*W*_, we can also examine their distributions, or (in the case of the polygenic trait) the distribution of the variances contributed by individual trait loci to the full population or within demes. Figures 4a and 4c show that theory for the infinite-island model accurately predicts the distribution of per-locus contributions to 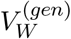 in simulations with *D* = 100 demes. However, infinite-island theory cannot predict the distribution of per-locus contributions to 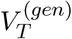, since (by definition) in the *D* → ∞ limit, the variance of *p* at any locus must be zero, and the distribution of any function of *p* must be a delta function. Thus, we investigate the effect of finite *D* on the distribution of per-locus contributions to 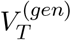 using only simulations. Figure S3 shows that, as expected, the distribution of variance contributions to 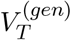 becomes narrower as the number of demes (and consequently, *N*_*tot*_) increases. Also, the distribution of variance contributions to 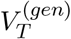 is comparably broad across loci of different effect sizes. Finally, we note that per-locus contributions to 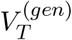 are much less broadly distributed than the corresponding contributions to 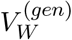 (compare Figs. 4a and 4c with Fig. S3 for *D* = 100). Thus, any locus is much more likely to contribute disproportionately to within-deme than total genic variance.

#### Finite number of trait loci

In the large *L* limit, the distribution of trait values is expected to be approximately normal. This assumption of normality underlies various QG predictions in the main text including eqs. (14), (15), (16) and (17). However, when selection per locus is strong, then the number of polymorphic loci can be rather small even if *L* is not too small. In this case, we expect the assumption of normally distributed trait values to break down, resulting in deviations between simulation results and theoretical predictions for the polygenic trait. This is also what we see in the Figure 2 for traits influenced by equal-effect loci of relatively large effect (*Ns* = 5). In these figures, simulation results deviate significantly from LD predictions for both low and high values of *Nm* (when overall levels of variation are low) and are, in fact, closer to the LE prediction.

**Figure S3.**
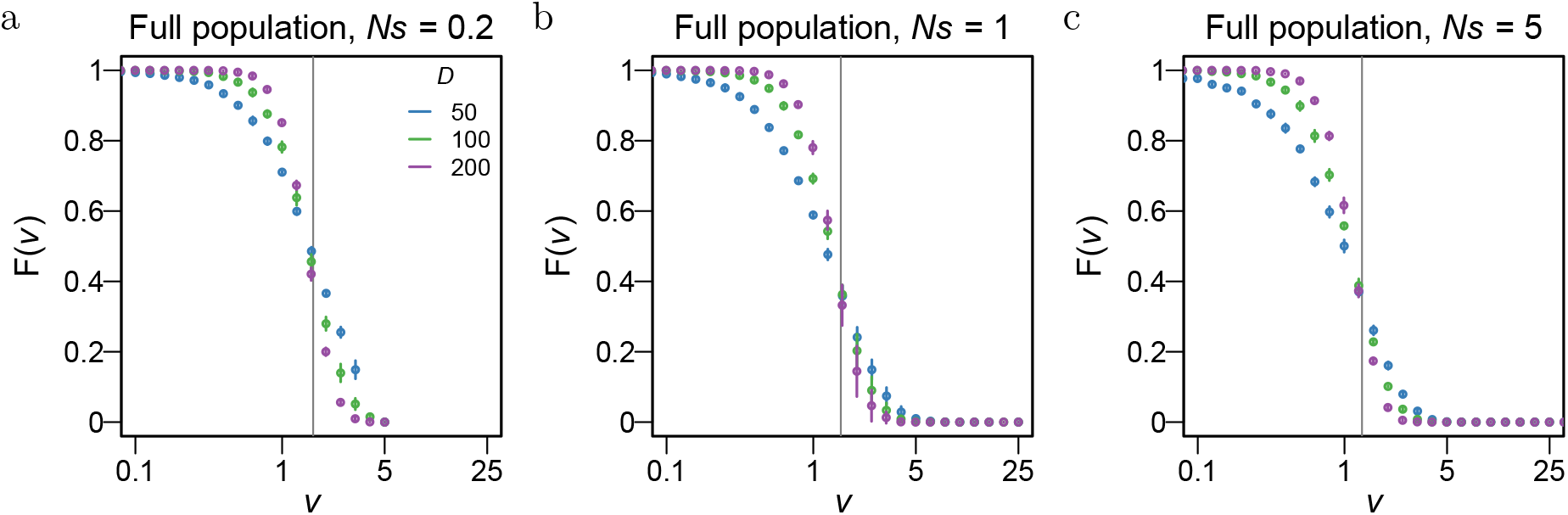
Cumulative probability *F* (*v*) that the contribution of a locus to 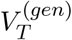 exceeds *v*, as a function of *v* (which is measured in units of *v*_*\**_), for a trait influenced by *L* = 200 loci of equal effect with scaled effect size per locus (a) *Ns* = 0.1 (b) *Ns* = 1.0 and (c) *Ns* = 5. Plots are obtained from individual-based simulations with different numbers of demes *D* (different colors), with other parameters being *N* = 200, *Nm* = 2 and *Nµ* = 0.01. The vertical line shows the expected variance contribution in the *D* → ∞ limit of the model (as obtained from LD predictions). As *D* increases, the cumulative distribution falls off more and more sharply, and is expected to approach a step function in the limit of very large *D*.

To better understand how the number of loci, *L*, influences the match between simulations and theory, we focus on a simple scenario of stabilizing selection on a trait influenced by equal-effect loci in a *single* panmictic population. In this case, the LE prediction for the (scaled) genic variance depend only on two scaled parameters: *Nµ* and *Ns* (and is independent of *L*), while LD predictions depend additionally on 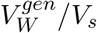 (which is 4*µL* times the scaled genic variance). This means that LD predictions for the (scaled) genic variance depend on three parameters: *Nµ, Ns* and *µL*. Thus, to test if simulation results approach LD predictions as *L* is increased, these three composite parameters must be held constant. This is accomplished by varying parameters according to: *L* → *xL, N* → *xN, µ* → *µ/x*, and *V*_*s*_ → *V*_*s*_*x* (where *x* is an arbitrary scaling factor), such that 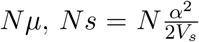 and *µL* remain unchanged. Note that under this kind of parameter scaling, increasing *L* causes 2*NµL* (which is the population-wide rate of trait-affecting mutations) to increase by the same factor.

Figure S4 shows how the scaled genic variance of a trait with *Ns* = 5, *Nµ* = 0.01 and *µL* = 0.01 depends on *NµL*. As in Fig. 2, genic variance in simulations is significantly lower than the LD prediction for small values of *NµL*. However, increasing *NµL* (by increasing *L* using the parameter scaling described above) causes simulation results to converge towards the LD prediction.

**Figure S4.**
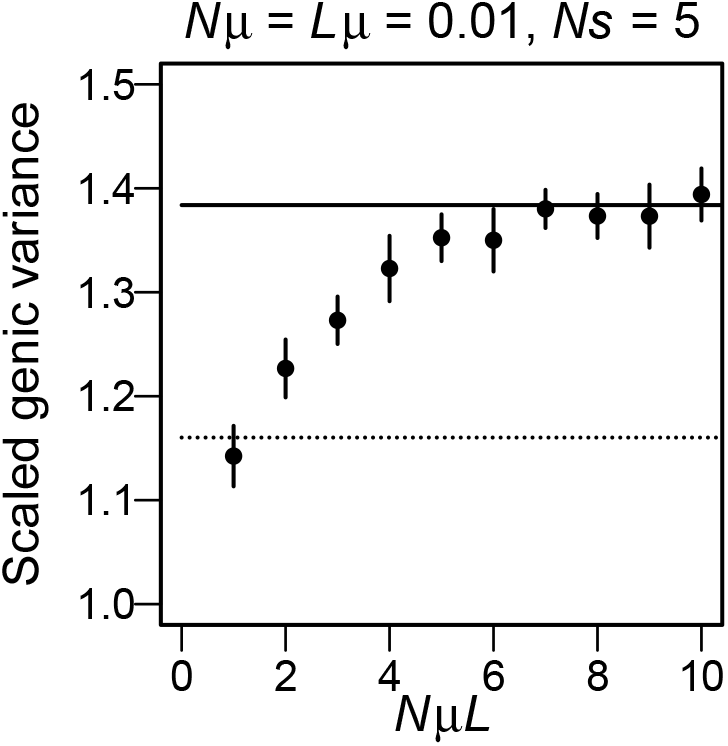
Scaled genic variance in a single panmictic population under stabilizing selection as a function of *NµL*, for *Nµ* = 0.01, *Ns* = 5 and *Lµ* = 0.01. Symbols show results of individual-based simulations; the dashed line shows the LE prediction; the solid line shows the prediction that accounts for the effects of LD on genic variance via *s*_eff_. To vary *NµL*, we increase *N*, simultaneously decreasing *µ* and increasing *L* and *V*_*s*_ by the same factor, such that *Nµ, Lµ* and 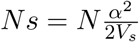remain unchanged. Simulation results approach the LD prediction at higher values of *NµL*. However, for smaller *NµL*, there is a discrepancy between the two due to there being too few polymorphic trait loci.

### SI.d The approach to equilibrium

In the main paper, we focus on equilibrium predictions for genic variance. However, populations may equilibrate very slowly under conditions of low gene flow, especially in scenarios where equilibration involves the crossing of fitness valleys by individual demes (that may initially occupy different adaptive peaks). To get some intuition for how the initial distribution of genotypes in a subdivided population influences equilibration times, we simulate a single underdominant locus as well as the full polygenic trait starting with two contrasting initial conditions IC1 and IC2 (described below) and under different levels of migration (Fig. S5).

**Figure S5.**
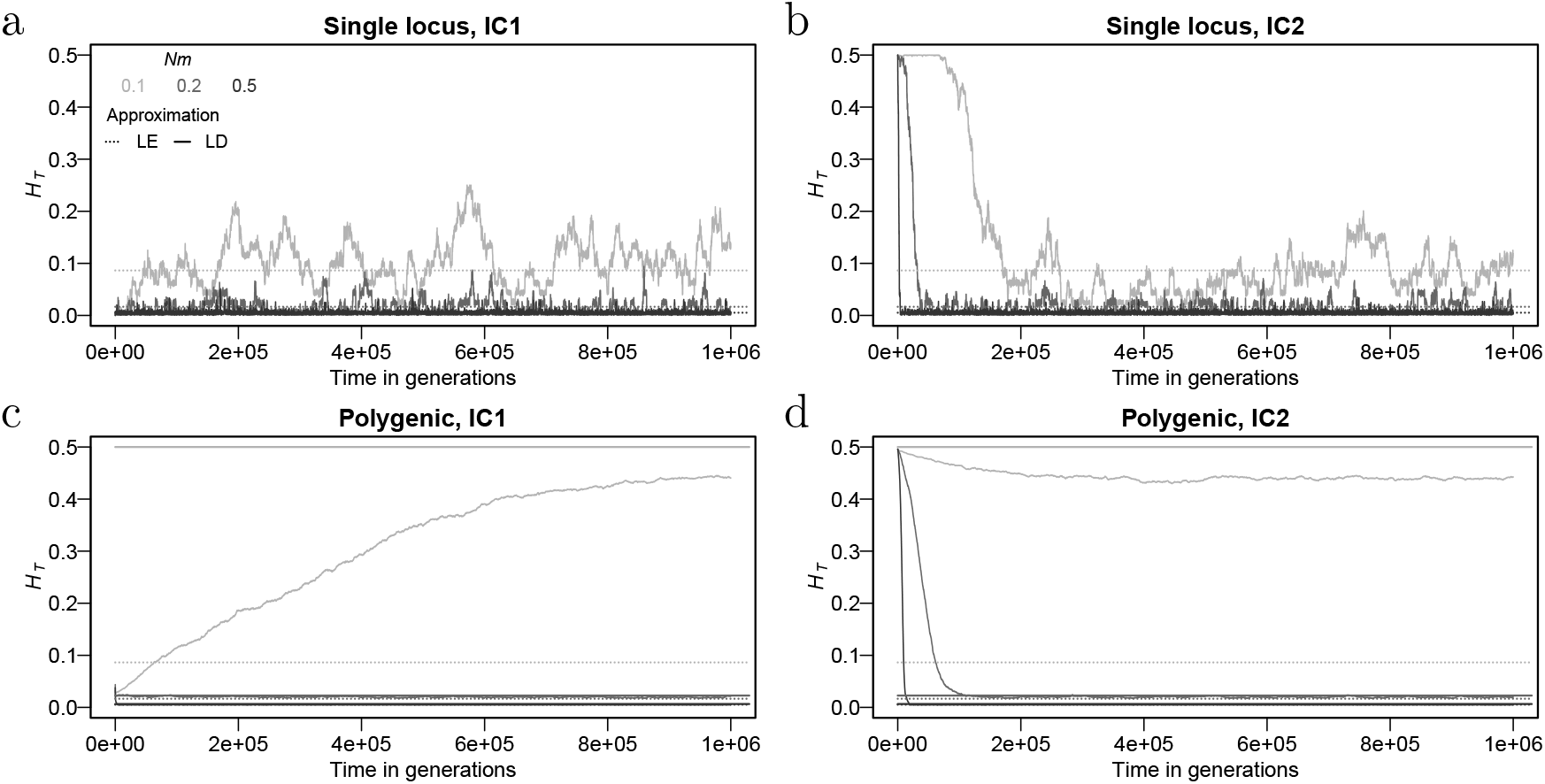
Total heterozygosity 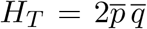as a function of time for: (a),(b) a single under-dominant locus; (c),(d) *L* = 200 equal-effect loci influencing a trait under stabilizing selection (where heterozygosities are averaged across all loci), for three different values of *Nm*. Plots show results of a single simulation run with *D* = 100, *N* = 200, *Nµ* = 0.01 and *Ns* = 5, that are initialised in one of two ways. In plots (a) and (c), the mean allele frequency *p* at any locus is drawn from a Beta distribution *B*[4*Nµ*, 4*Nµ*]; the allele frequency in each deme is then set to this value (IC1). In plots (b) and (d), allele frequencies in each deme are drawn independently from a Beta distribution *B*[4*Nµ*, 4*Nµ*] (IC2). The two initial conditions IC1 and IC2 thus correspond respectively to very low or maximal *H*_*T*_ at the start of the simulation. Dashed lines represent the theoretical predictions for *H*_*T*_ at equilibrium. Equilibration takes longer at lower levels of migration. Further, equilibration is slower when it involves a large change in *H*_*T*_ from very low to high values (or vice versa), as this necessitates the crossing of fitness valleys (generated by underdominant selection) at multiple loci.

Under the first kind of initial condition (IC1), we start with all demes having the same allele frequency at a given locus, while under IC2, allele frequencies across different demes are sampled independently from a Beta distribution (details in caption of Fig. S5). In essence, all demes either initially occupy the same adaptive peak (IC1) or are uniformly distributed across all possible adaptive peaks (IC2). Thus, the total heterozygosity across the population as a whole is either extremely low (IC1) or at its maximum possible value of 1*/*2 (IC2).

As expected, equilibration is slower for smaller values of *Nm* (Fig. S5). Further, equilibration is also slower when it involves a qualitative change in the state of the population from its initial state. For example, equilibration takes longer when a population must evolve from initially high *H*_*T*_ (IC2; different alleles common in different demes) to low *H*_*T*_ (where the same allele is common in all demes) than when it starts already at low *H*_*T*_ : compare, for example, IC1 vs. IC2 for *Nm* = 0.2 in Figs. S5c and S5d.

### SI.e Within-deme and total genetic variance

Figures 2e and 2d in the main paper show how the ratio of the genetic to genic variance changes with the strength of selection and migration. This ratio is less than 1 (especially for strong selection and weak migration), as there is negative LD between trait-increasing (or trait-decreasing) alleles within demes and at the level of the full population. Thus, 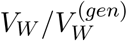 and 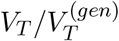can be used as proxies for the total amount of LD between trait loci within demes and across the full population respectively.

Here, we examine how the genetic variances *V*_*W*_ and *V*_*T*_ themselves (rather than scaled by genic variance) depend on various parameters for a trait with equal-effect loci. Figure S6 shows that both the within-deme and total genetic variances are maximized at intermediate levels of migration (*Nm* ≃ *Nm*_crit_). Further, while strong population structure inflates total genic variance by orders of magnitude (Fig. 2a), especially for traits influenced by many large-effect loci, its effect on the total genetic variance is quite modest. This is consistent with the observation that 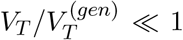 at low migration rates (Fig. 2d). Finally, we note that *V*_*T*_ ≃ *V*_*W*_, except for the case of *Ns* = 0.1 (and *Nm* ≪ 1). This is consistent with the rather low values of *Q*_*ST*_ observed in Fig. 2f and reflects the fact that very little of the total genetic variance in a population is due to among-deme differences in trait means, since these are quite tightly constrained under stabilizing selection.

**Figure S6.**
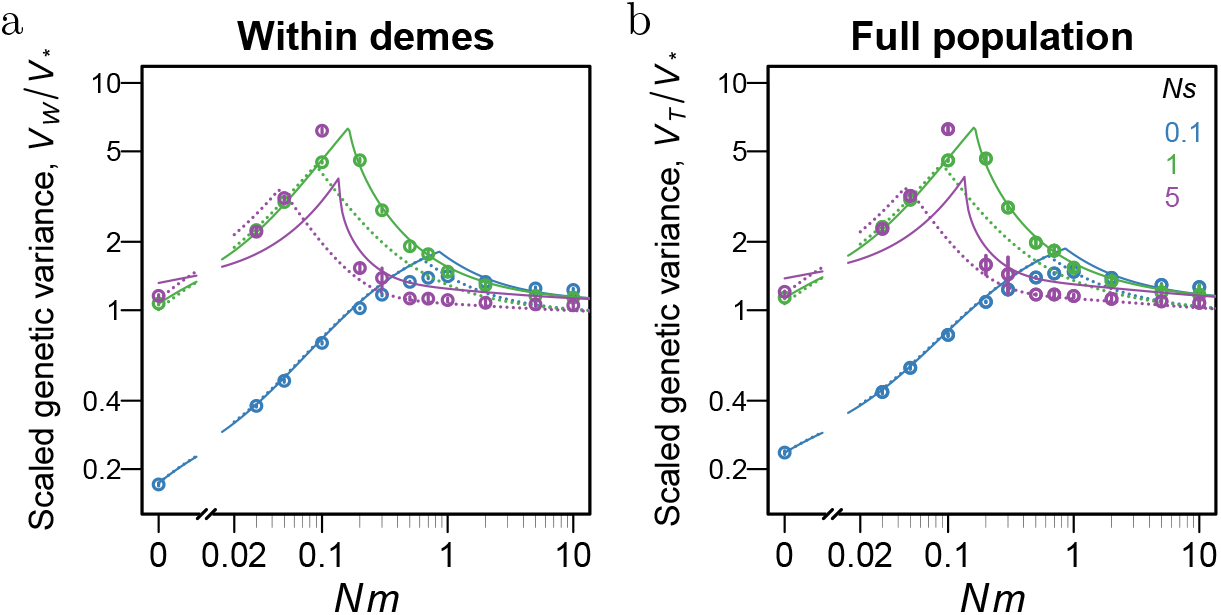
(a) Within-deme genetic variance *V*_*W*_ */V*_*\**_ and (b) total genetic variance *V*_*T*_ */V*_*\**_ as a function of *Nm* for a trait influenced by equal-effect loci, for 3 different values of *Ns* (scaled effect size per locus). All parameters are the same as in Fig. 2 (*D* = 100, *N* = 200, *L* = 200 and *Nµ* = 0.01), and symbols show results from the same set of individual-based simulations. Theoretical predictions are based on eqs. (14), (19) and (21), together with single-locus theory for the genic variances – either with or without effective parameters (solid vs. dashed lines).

### SI.f Dependence of various quantities on mutation rate per locus

In the main paper, we always keep the scaled mutation rate fixed at *Nµ* = 0.01. Here, we explore the dependence of various quantities on mutation rate based on our theoretical predictions. Figures S7a,b show that the scaled genic variances depend primarily on the ratio *µ/s*, and not on *Nµ* and *Ns* separately. Similarly the distribution of per-locus contributions to 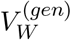 and the proportion of 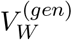 accounted for by loci with variance exceeding a certain value (see also Figures 4a, 4b) also depend primarily on *µ/s* (Figures S7c,d).

**Figure S7.**
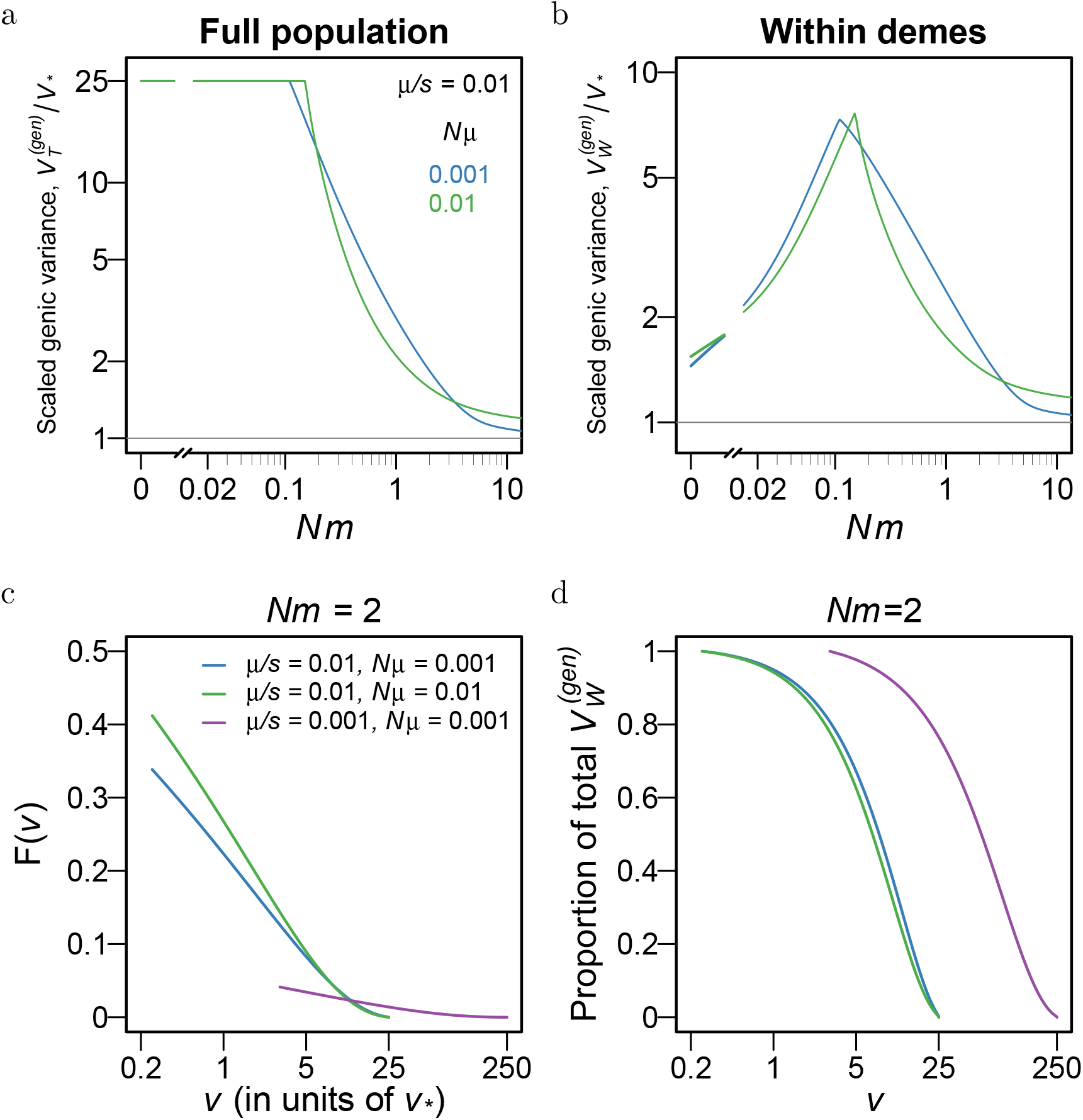
(a),(b): The expected (a) total genic variance across the full population, 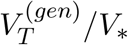, and (b) within-deme genic variance, 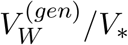, as a function of *Nm*, for a trait influenced by *L* = 200 equal-effect loci. Genic variances are scaled by *V*_*\**_ = 4*µLV*_*s*_. The different colors show theoretical LD predictions for different values of *Ns* and *Nµ* (with *µ/s* = 0.01 held fixed). (c),(d): (c) Cumulative probability *F* (*v*) that the variance contributed by a locus exceeds *v*, and (d) proportion of within-deme genic variance 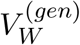 explained by loci with variance exceeding *v*, as a function of *v*, for a trait influenced by *L* = 200 loci of equal effect, for *Nm* = 2. All per-locus variances are measured in units of *v*_*\**_ = 4*µV*_*s*_. The different colors show theoretical LD predictions for different values of *Nµ*, keeping either *µ/s* or *Ns* fixed.

## References

Barghi, N., Hermisson, J., and Schlötterer, C. (2020). Polygenic adaptation: a unifying framework to understand positive selection. Nature Reviews Genetics, 21(12):769–781.

Barghi, N., Tobler, R., Nolte, V., Jakšić, A. M., Mallard, F., Otte, K. A., Dolezal, M., Taus, T., Kofler, R., and Schlötterer, C. (2019). Genetic redundancy fuels polygenic adaptation in drosophila. PLOS Biology, 17(2):1–31.

Barton, N. H. (1999). Clines in polygenic traits. Genetical Research, 74(3):223–236.

Barton, N. H. and Keightley, P. D. (2002). Understanding quantitative genetic variation. Nature Reviews Genetics, 3(1):11–21.

Barton, N. H. and Rouhani, S. (1993). Adaptation and the ‘shifting balance’. Genetics Research, 61(1):57–74.

Battey, C. J., Ralph, P. L., and Kern, A. D. (2020). Space is the place: Effects of continuous spatial structure on analysis of population genetic data. Genetics, 215(1):193–214.

Berg, J. J., Li, X., Riall, K., Hayward, L. K., and Sella, G. (2025). Mutation-selection-drift balance models of complex diseases. Genetics, 231(4):iyaf220.

Bertram, J. and Shafiei, Z. (2025). Strong amplification of quantitative genetic variation under a balance between mutation and fluctuating stabilizing selection. bioRxiv.

Booker, T. R., Yeaman, S., and Whitlock, M. C. (2021). Global adaptation complicates the interpretation of genome scans for local adaptation. Evolution Letters, 5(1):4–15.

Bulmer, M. (1971). The effect of selection on genetic variability. The American Naturalist, 105(943):201–211.

Bulmer, M. (1972). The genetic variability of polygenic characters under optimizing selection, mutation and drift. Genetics Research, 19(1):17–25.

Bürger, R. and Gimelfarb, A. (2002). Fluctuating environments and the role of mutation in maintaining quantitative genetic variation. Genetical Research, 80(1):31–46.

Bürger, R., Wagner, G. P., and Stettinger, F. (1989). How much heritable variation can be maintained in finite populations by mutation-selection balance? Evolution, 43(8):1748–1766.

Charlesworth, B. (2013). Stabilizing selection, purifying selection, and mutational bias in finite populations. Genetics, 194(4):955–971.

Duncan, L., Shen, H., Gelaye, B., Meijsen, J., Ressler, K., Feldman, M., Peterson, R., and Domingue, B. (2019). Analysis of polygenic risk score usage and performance in diverse human populations. Nature communications, 10(1):3328–9.

Goldstein, D. B. and Holsinger, K. E. (1992). Maintenance of polygenic variation in spatially structured populations: Roles for local mating and genetic redundancy. Evolution, 46(2):412–429.

Guillaume, F. and Whitlock, M. C. (2007). Effects of migration on the genetic covariance matrix. Evolution, 61(10):2398–2409.

Höllinger, I., Wölfl, B., and Hermisson, J. (2023). A theory of oligogenic adaptation of a quantitative trait. Genetics, 225(2):iyad139.

Jouganous, J., Long, W., Ragsdale, A. P., and Gravel, S. (2017). Inferring the joint demographic history of multiple populations: Beyond the diffusion approximation. Genetics, 206(3):1549– 1567.

Kimura, M. (1964). Diffusion models in population genetics. Journal of Applied Probability, 1(2):177–232.

Kimura, M. (1965). Attainment of quasi linkage equilibrium when gene frequencies are changing by natural selection. Genetics, 52(5):875–890.

Kobayashi, Y., Hammerstein, P., and Telschow, A. (2008). The neutral effective migration rate in a mainland-island context. Theoretical Population Biology, 74(1):84–92.

Koch, E., Connally, N. J., Baya, N., Reeve, M. P., Daly, M., Neale, B., Lander, E. S., Bloemendal, A., and Sunyaev, S. (2024). Genetic association data are broadly consistent with stabilizing selection shaping human common diseases and traits. bioRxiv.

Lande, R. (1975). The maintenance of genetic variability by mutation in a polygenic character with linked loci. Genetics Research, 26(3):221–235.

Lande, R. (1976). Natural selection and random genetic drift in phenotypic evolution. Evolution, pages 314–334.

Lande, R. (1991). Isolation by distance in a quantitative trait. Genetics, 128(2):443–452.

Latta, R. (1998). Differentiation of allelic frequencies at quantitative trait loci affecting locally adaptive traits. The American Naturalist, 151(3):283–292. PMID: 18811359.

Latter, B. (1960). Natural selection for an intermediate optimum. Australian Journal of Biological Sciences, 13(1):30–35.

Le Corre, V. and Kremer, A. (2003). Genetic variability at neutral markers, quantitative trait loci and trait in a subdivided population under selection. Genetics, 164(3):1205–1219.

Le Corre, V. and Kremer, A. (2012). The genetic differentiation at quantitative trait loci under local adaptation. Molecular Ecology, 21(7):1548–1566.

Leinonen, T., McCairns, R. S., O’hara, R. B., and Merilä J. (2013). Qst-fst comparisons: evolutionary and ecological insights from genomic heterogeneity. Nature Reviews Genetics, 14(3):179–190.

Lythgoe, K. A. (1997). Consequences of gene flow in spatially structured populations. Genetical Research, 69(1):49–60.

MacPherson, A., Hohenlohe, P., and Nuismer, S. (2015). Trait dimensionality explains widespread variation in local adaptation. Proc Biol Sci., 282(1802):20141570.

MacPherson, A. and Nuismer, S. L. (2017). The probability of parallel genetic evolution from standing genetic variation. Journal of Evolutionary Biology, 30(2):326–337.

McDonald, T. K. and Yeaman, S. (2018). Effect of migration and environmental heterogeneity on the maintenance of quantitative genetic variation: a simulation study. Journal of Evolutionary Biology, 31(9):1386–1399.

Morjan, C. L. and Rieseberg, L. H. (2004). How species evolve collectively: implications of gene flow and selection for the spread of advantageous alleles. Molecular Ecology, 13(6):1341–1356.

Nagylaki, T. (1978). Random genetic drift in a cline. Proceedings of the National Academy of Sciences, 75(1):423–426.

Negm, S. and Veller, C. (2024). The effect of long-range linkage disequilibrium on allele-frequency dynamics under stabilizing selection. bioRxiv, pages 2024–06.

Patel, R. A., Weiß, C. L., Zhu, H., Mostafavi, H., Simons, Y. B., Spence, J. P., and Pritchard, J. K. (2024). Characterizing selection on complex traits through conditional frequency spectra. Genetics, 229(4):iyae210.

Phillips, P. C. (1996). Maintenance of polygenic variation via a migration–selection balance under uniform selection. Evolution, 50(3):1334–1339.

Ragsdale, A. P. (2025). Archaic introgression and the distribution of shared variation under stabilizing selection. PLOS Genetics, 21(3):1–20.

Ralph, P. L. and Coop, G. (2015). The role of standing variation in geographic convergent adaptation. The American Naturalist, 186(S1):S5–S23.

Robertson, A. (1956). The effect of selection against extreme deviants based on deviation or on homozygosis. Journal of Genetics, 54:236–248.

Sachdeva, H. (2022). Reproductive isolation via polygenic local adaptation in sub-divided populations: Effect of linkage disequilibria and drift. PLOS Genetics, 18(9):e1010297.

Sella, G. and Barton, N. H. (2019). Thinking about the evolution of complex traits in the era of genome-wide association studies. Annual review of genomics and human genetics, 20:461–493.

Simons, Y. B., Bullaughey, K., Hudson, R. R., and Sella, G. (2018). A population genetic interpretation of gwas findings for human quantitative traits. PLoS biology, 16(3):e2002985.

Simons, Y. B., Mostafavi, H., Zhu, H., Smith, C. J., Pritchard, J. K., and Sella, G. (2025). Simple scaling laws control the genetic architectures of human complex traits. PLOS Biology, 23(10):1–21.

Slatkin, M. (1978). Spatial patterns in the distributions of polygenic characters. Journal of Theoretical Biology, 70(2):213–228.

Spence, J., Mostafavi, H., Ota, M., Milind, N., Gjorgjieva, T., Smith, C. J., Simons, Y. B., Sella, G., and Pritchard, J. K. (2026). Specificity, length and luck drive gene rankings in association studies. Nature, 649(8098):918–925.

Spitze, K. (1993). Population structure in daphnia obtusa: quantitative genetic and allozymic variation. Genetics, 135(2):367–374.

Steiner, M. C., Rice, D. P., Biddanda, A., Ianni-Ravn, M. K., Porras, C., and Novembre, J. (2025). Study design and the sampling of deleterious rare variants in biobank-scale datasets. Proceedings of the National Academy of Sciences, 122(23):e2425196122.

Surendranadh, P. and Sachdeva, H. (2025). Effect of assortative mating and sexual selection on polygenic barriers to gene flow. Evolution, 79(7):1185–1198.

Tufto, J. (2000). Quantitative genetic models for the balance between migration and stabilizing selection. Genetical Research, 76(3):285–293.

Turelli, M. (1984). Heritable genetic variation via mutation-selection balance: Lerch’s zeta meets the abdominal bristle. Theoretical population biology, 25(2):138–193.

Veller, C. and Coop, G. M. (2024). Interpreting population-and family-based genome-wide association studies in the presence of confounding. PLoS Biology, 22(4):e3002511.

Veller, C. and Simons, Y. (2024). Stabilizing selection generates selection against introgressed dna. bioRxiv.

Walsh, B. and Lynch, M. (2018). Maintenance of quantitative genetic variation. In Evolution and Selection of Quantitative Traits. Oxford University Press.

Wang, Y., Guo, J., Ni, G., Yang, J., Visscher, P. M., and Yengo, L. (2020). Theoretical and empirical quantification of the accuracy of polygenic scores in ancestry divergent populations. Nature communications, 11(1):3865.

Weir, B. S. and Cockerham, C. C. (1984). Estimating f-statistics for the analysis of population structure. Evolution, 38(6):1358–1370.

Westram, A. M., Stankowski, S., Surendranadh, P., and Barton, N. (2022). What is reproductive isolation? Journal of Evolutionary Biology, 35(9):1143–1164.

Whitlock, M. C. (1999). Neutral additive genetic variance in a metapopulation. Genetical Research, 74(3):215–221.

Whitlock, M. C. (2008). Evolutionary inference from qst. Molecular Ecology, 17(8):1885–1896.

Wright, S. (1931). Evolution in mendelian populations. Genetics, 16(2):97–159.

Wright, S. (1935). Evolution in populations in approximate equilibrium. Journal of Genetics, 30(2):257–266.

Wright, S. (1937). The distribution of gene frequencies in populations. Proceedings of the National Academy of Sciences, 23(6):307–320.

Wright, S. (1949). The genetical structure of populations. Annals of Eugenics, 15(1):323–354.

Yair, S. and Coop, G. (2022). Population differentiation of polygenic score predictions under stabilizing selection. Philosophical Transactions of the Royal Society B, 377(1852):20200416.

Yeaman, S., Gerstein, A. C., Hodgins, K. A., and Whitlock, M. C. (2018). Quantifying how constraints limit the diversity of viable routes to adaptation. PLoS genetics, 14(10):e1007717.

